# An intercellular signaling pathway in the mouse retina connects Kv2.1, GLT-1, and nitric oxide synthase 1 to optic nerve regeneration

**DOI:** 10.1101/2025.05.07.652294

**Authors:** Hui-ya Gilbert, Yiqing Li, Kenya Yuki, Kumiko Omura, Melody Walter, Nicholas Hanovice, Yuqin Yin, Paul Huang, Niels C. Danbolt, Zhe Liu, Jiahui Tang, Qi Zhang, Elias Aizenman, Stephen J. Lippard, Larry I. Benowitz, Paul A. Rosenberg

## Abstract

We report here that a multicellular signaling pathway in the mouse retina targeting glutamate homeostasis and nitric oxide production is activated upon optic nerve injury and modulates retinal ganglion cells’ (RGCs’) ability to mount a robust regenerative response. A novel highly sensitive and specific NO sensor (FL2) revealed that optic nerve injury leads to a rapid, prolonged elevation of NO in the inner retina. Amacrine cell-specific genetic deletion of the neuron-specific isoform of nitric oxide synthase (NOS1, nNOS) or the NOS1 inhibitor N(w)-propyl-L-arginine (L-NPA) suppressed optic nerve regeneration. Steps leading to NOS1 activation are shown to include Kv2.1 phosphorylation and activation, reversal of glutamate uptake by the glutamate transporter GLT-1 (EAAT2), and subsequent NMDA receptor activation. Conditional knockout of the glutamate transporter GLT-1 in bipolar cells, intraocular injection of the GLT-1 blockers dihydrokainate (DHK) or WAY213613, the N-methyl-D-aspartate (NMDA) receptor antagonist D-2-amino-5-phosphonovalerate (D-AP5), or the Kv2.1 blockers RY796 or stromatoxin all suppressed NO generation and strongly diminished RGCs’ ability to respond to Pten deletion and other factors. Thus, optic nerve injury activates a sequence of pathophysiological changes in retinal interneurons that gates RGCs’ ability to regenerate injured axons.

**Significance Statement:** Cells use extracellular signals to coordinate responses to injury and to promote repair and regeneration. The NMDA receptor, which has a high affinity for glutamate and is functionally coupled to nitric oxide synthase, has the potential to mediate such responses by responding to changes in extracellular glutamate. In the mouse retina after optic nerve injury, potassium channels in retinal ganglion cells, glutamate transporters in neighboring cells, and NMDA receptors in amacrine cells, are all involved in a pathway that regulates the generation of nitric oxide, which we show promotes axon regeneration after injury.

---

Following optic nerve injury (ONI), mature retinal ganglion cells (RGCs) cannot regrow axons and soon begin to die, precluding visual recovery (1, 2). Cell-intrinsic factors known to influence RGC survival and axon regeneration include regulators of signal transduction (3–10) and apoptosis (11–15), transcription factors (16–20), and epigenetic regulators (21–24), among others (25–28). RGC survival and optic nerve regeneration are also influenced by cell non-autonomous factors that include myelin debris (29, 30), scar-forming cells (31–33), inflammatory cells (34–41), and, as recently discovered, amacrine cells (ACs), the inhibitory interneurons of the retina (42, 43). Of the many types of ACs that have been identified, sparse subsets expressing nitric oxide synthase-1 (NOS1) are seen within the inner nuclear and ganglion cell layers (INL, GCL) (44–48). NO is elevated after ONI and is reported to negatively affect RGC survival in mammals (49–51) and suppress axon regeneration in Drosophila (52), whereas positive effects of NO on axon regeneration have been reported in goldfish (53). The mechanisms underlying NO elevation in the retina after ONI have not been characterized.

NMDA receptors constitute a major molecular signaling hub in neurons and, unlike non-NMDA receptors, have a micromolar affinity for the excitatory neurotransmitter glutamate, implying the need for tight regulation of glutamate homeostasis (54, 55). Opening of NMDA receptor-gated channels allows an influx of Ca^2+^, which interacts with intracellular calmodulin to activate NOS1 and generate NO (56–58). Several studies suggest that NMDA receptors contribute to RGC loss after optic nerve injury (59, 60), but the possible relationship of NMDA receptor activation to NO generation and to optic nerve regeneration is unknown. Further, although it is generally assumed that NOS1 is activated by synaptically released glutamate (61–63), an alternative mechanism after optic nerve injury could be dysregulation of extracellular glutamate homeostasis mediated by reversal of glutamate transport (64–69) resulting in activation of high-affinity NMDA-type glutamate receptors (70–73). Whereas NO has been reported to regulate glutamate metabolism and glutamate transporter function (74), there is no prior evidence that alterations in glutamate transporter function regulate nitric oxide generation.

GLT-1 is the major glutamate transporter in the forebrain (75, 76) where it is expressed primarily in astrocytes (77–79) and in excitatory axon terminals (80–83). In the retina, GLT-1 has been reported to be associated with a population of amacrine cells, probably GABAergic (84), cone photoreceptors, and subpopulations of bipolar cells (84–86). Glutamate transporters are secondary active transporters that use the energy stored in ionic gradients across the plasma membrane to perform concentration work. There has been substantial interest in the mechanisms that underlie reversal of glutamate transport and efflux of glutamate into the extracellular space resulting in excitotoxic injury (67, 87–90). Recent evidence suggests that alterations in the extracellular ionic milieu due to neuronal activity (91–96), resulting in a decrease in extracellular sodium or increase in extracellular potassium, could significantly alter the function of glutamate transporters.

The potassium channel Kv2.1 is a major component of the delayed rectifier current in neurons (97, 98), and has a unique anatomical and functional relationship with the glutamate transporter GLT-1. Clusters of Kv2.1 in regions of the neuronal plasma membrane in contact with the endoplasmic reticulum are in apposition to regions of high density of expression of GLT-1 in astrocytic plasma membranes (99). Importantly, Kv2.1 has been shown to regulate potassium flux across the plasma membrane and is critical in initiating apoptosis by promoting activation of enzymes inhibited at normal intracellular concentrations of potassium (100–104). The Kv2.1-GLT-1 complex has been postulated as an “injury sensor” monitoring neuronal stress and glutamate dyshomeostasis by responding via activation of high affinity NMDA receptors, Kv2.1 declustering, and a hyperpolarizing shift in the voltage-dependence of Kv2.1 channels (105, 106). A role for Kv2.1 operating in the opposite direction, whereby changes in Kv2.1 expression or activation on the plasma membrane, perturbation of extracellular potassium and related alteration of GLT-1 function leading to NMDA receptor activation, has not previously been considered. We report here evidence for such a pathway that induces NMDA receptor activation in NOS1-positive ACs, resulting in a rapid and persistent elevation of NO that enables RGCs to respond robustly to a pro-regenerative therapy.

## Results

### Rapid and prolonged NO elevation after optic nerve injury

To investigate retinal NO dynamics following optic nerve injury, our initial experiments used both the membrane impermeable, non-trappable fluorescent NO probe 2-(6-hydroxy-4,5-bis[(2-methylquinolin-8-ylamino) methyl]-3-oxo-3H-xanthen-9-yl)benzoic acid (FL2); and the acetylated, membrane-permeant, trappable variant, 2-(4,5-bis[(6-(2-ethoxy-2-oxoethoxy)-2-methylquinolin-8-ylamino)methyl]-6-hydroxy-3-oxo-3H-xanthen-9-yl)benzoic acid (FL2E) (107–109). The most important biological scavenger of nitric oxide is the superoxide anion O_2_^∸^, which reacts with NO in a diffusion-limited reaction to form peroxynitrite (110, 111). Because levels of the superoxide radical increase in RGCs following optic nerve crush (ONC) (112), the reaction of NO with superoxide could poten-tially diminish our ability to detect NO *per se* using either probe, as both are insensitive to peroxynitrite (107). We therefore investigated retinal NO signals in the absence and presence of copper-zinc superoxide dismutase (SOD, 2 U/eye) to scavenge O_2_^∸^. FL2 injected intravitreally 60 min prior to euthanasia revealed little fluorescent signal in freshly prepared sections of the normal retina, which increased slightly but significantly in the presence of SOD (Fig. 1A,B).

**Figure 1.**
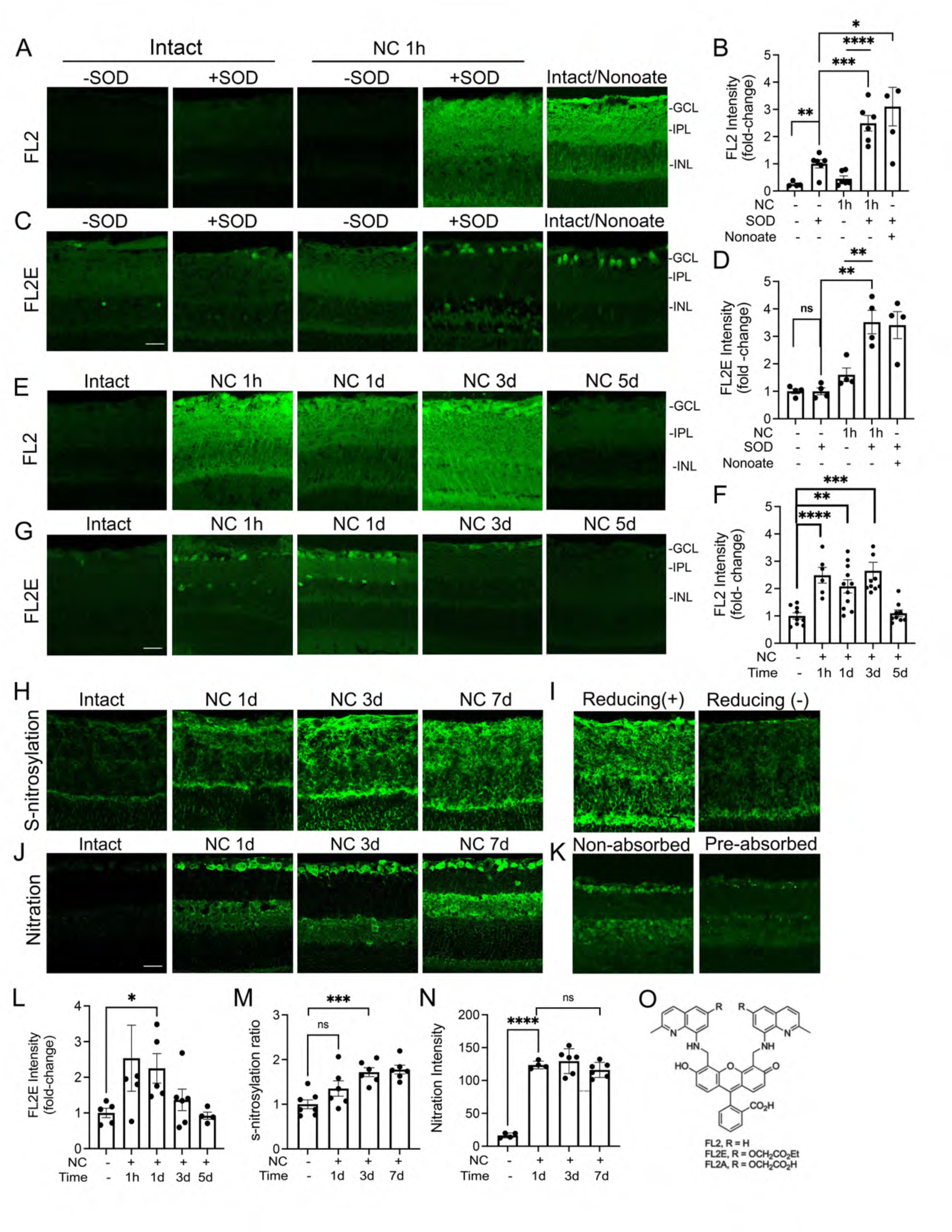
Rapid elevation of nitric oxide and its derivatives in the inner retina after optic nerve injury. (A,B) FL2, a specific detector of extracellular NO, and FL2E, a detector of intracellular NO (C,D), both reveal an SOD-dependent increase in a 520 nm NO signal after ONC. Superoxide dismutase (SOD) chelates superoxide radicals and prevents conversion of NO to peroxynitrite. Whereas little signal is seen with either detector in the intact retina with or without SOD, the presence of SOD allows visualization of elevated extra-and intracellular NO within an hour after ONC (A,C). As a positive control for the sensitivity of the detectors, intraocular injection of the NO donor DETA-NONOate in the intact retina elevates both extra-and intracellular NO signals (*green fluorescence*). Scale bar: 25 µM. (B,D) Quantitation of FL2 in A and FL2E in C. NO signal visualized with and without SOD in retinal sections (Mean ± SEM, n = 4-6 reti-nas/condition; \*\**P < 0.0046, t = 3.886*; \*\*\**P =* 0.001, t = 4.594. \*\*\*\**P <* 0.0001, *t* = 6.690 in B; ** *P =* 0.0013*, t =* 5.652 *(SOD only vs NC w/ SOD)*; \*\**P =* 0.0081*, t =* 3.890 *(NC w/SOD vs NC w/o SOD;* ns, *P =* 0.0604, *t* = 2.308 in D). (E-G) Time-course of NO elevation after ONC. Note early onset and long persistence of NO signal detected with both FL2 (E) and FL2E (G). Scale bar: 25 µM. (F) Quantitation of NO signal (FL2) in the inner retina 1 hour to 5 days after ONC (n = 4-12 mice/condition, \*\**P =* 0.023, t = 4.834 *(Intact vs NC 1hour)*; \*\**P =* 0.0011, t = 4.075 *(Intact vs NC 1 Day);* \*\*\**P =* 0.0004, *t* = 4.923). (H) Enduring changes in protein S-nitrosylation in the inner retina after ONC. Protein S-nitrosylation is detected using the biotin-switch assay at the indicated times after ONC. (I) Control for specificity of the S-nitrosylation signal shows an absence of reaction product in the presence of the labeling reagent but without reducing buffer. (J) Changes in protein nitration in the inner retina detected by anti-nitrotyrosine immunostaining at the indicated times after ONC. (K) Control for the specificity of protein nitration: pre-adsorption of the primary antibody with 3-nitrotyrosine. Scale bar, 25 µM. (L) as in F but for FL2E probe (n = 4 mice/condition, \**P =* 0.0365, *t* = 2.874). (M, N) Quantitation of S-nitrosylation in the inner plexiform layer (IPL, M) and nitration in the ganglion cell layer (N) over time (n = 5-6 retinas/condition; ns = 0.0913, *t* = 1.850; \*\*\**P =* 0.003, t = 5.101 in M; n = 4-6/condition; \*\*\*\**P* < 0.0001, *t* = 30.86; ns = 0.564, *t* = 0.6012). (O) Structure of FL sensors. The membrane-permeable (FL2E) and-impermeable (FL2) forms differ only at position R [redrawn from ref. (107)].

Within an hour after ONC, there was still no detectable NO signal in the absence of SOD (Fig. 1A, panel 3 and 1B). However, in the presence of SOD, the signal increased 2.5-fold over baseline (Fig. 1A, panel 4 vs. 2, Fig. 1B, \*\*\**P <* 0.001, t = 4.594) and 4-fold relative to the post-crush NO signal measured without SOD1 (Fig. 1A, panel 4 vs. 3; Fig. 1B, \*\*\*\**P <* 0.0001, t = 6.690). As a positive control, injection of the NO donor diethylenetriamine NONOate (DETA-NONOate) (10 µM) in normal eyes resulted in a ∼ 3-fold increase in the fluorescent signal in the inner retina (Figs. 1A panels 5 vs. 2; Fig. 1B, \*\**P <* 0.0111, t = 3.183), verifying the ability of the sensor to effectively detect changes in NO levels. The esterified form of the sensor, FL2E, is membrane-permeable and revealed a qualitatively different pattern of staining from FL2, dominated by intracellular staining in the GCL (Fig. 1C). However, despite the qualitative differences in staining patterns, the changes found with FL2E were quantitatively similar to those obtained with FL2, that is, a 3-fold increase in signal intensity relative to the normal control retina within an hour after ONC (Fig. 1C, panels 4 vs. 2; Fig.1D \*\**P* = 0.0013, t = 5.652), dependence of the signal on SOD1 (Fig. 1C, panels 4 vs. 3; Fig.1D \*\**P* = 0.0081, t = 3.890), and a similar effect of DETA-NONOate (Fig. 1C, panel 5, Fig. 1D). Investigating the time-course of NO elevation, we used the same two sensors (plus SOD) at time points ranging from 1 h to 7 days post-optic nerve injury, with sensors introduced into the eye 1 h before euthanasia in all cases. Both sen-sors again showed an elevated NO signal within the first hour post-ONC that persisted up to 3 days in the case of FL2 but that declined in the case of FL2E between 1 and 3 days (Fig. 1E-F; Fig. 1G, L).

Because nitric oxide can react with free thiol groups in proteins, we investigated whether optic nerve injury leads to protein S-nitrosylation, a post-translational modification that can regulate protein activity, localization, and stability (113–116). Using the biotin-switch reaction to reveal sites of S-nitrosylation (117), we found that optic nerve injury resulted in a gradual increase in protein S-nitrosylation (Fig. 1H, panels 2 - 4 vs. 1), attaining statistical significance by 3 days (Fig. 1M,\*\*\**P* = 0.003, t = 5.101) and remaining elevated for at least 7 days. The specificity of the staining was demonstrated by its loss when the procedure was carried out in the absence of the reducing buffer that converts SNO groups to thiols via transnitrosation with ascorbate (Fig. 1I).

Conversion of NO to peroxynitrite by reaction with superoxide, as demonstrated by the effects of SOD on fluorescence intensity of FL2 and FL2E staining, appears to be a major route for the metabolism of NO in the retina after ON crush. Peroxynitrite can react with protein tyrosine residues to generate 3-nitrotyrosine, a modification that may alter protein function (118, 119).

Immunostaining revealed a rapid and persistent elevation of 3-nitrotyrosine in the GCL and inner nuclear layer (INL) that endured for at least 7 days after optic nerve damage (Fig. 1J), As a control for the specificity of the immunostaining, we found that the signal was lost when the primary antibody was pre-absorbed with 3-nitrotyrosine (Fig. 1K).

### NO Synthase-1 is the primary source of NO after optic nerve injury

Our initial approach to identify the source of NO used two inhibitors that target different NOS isoforms. L-Nω-propyl-L-arginine (L-NPA) is a potent, selective inhibitor of NO synthase-1 [NOS1, neuronal NOS (nNOS); *Ki* NOS1 = 57 nM], with a 150-fold lower affinity to NOS3 [endothelial NOS (eNOS); *Ki* = 8.5 μM], and a > 3000-fold lower (*Ki* =180 μM) (120, 121) affinity for NOS2 [inducible NOS (iNOS)]. On the other hand, 1400W is highly selective as a NOS2 inhibitor compared with its potency as an inhibitor of NOS3 and NOS1 (5000-fold and 250-fold, respectively) (122, 123). At a concentration of 100 µM, L-NPA reduced the NO signal after optic nerve injury (decreasing the difference between baseline and optic nerve injury by ∼ 2/3: Fig. 2A,B, \**P* = 0.0145, t = 2.634), whereas 1400W had a lesser, non-significant, effect (Fig. 2A, B, *P* = 0.1008, t = 1.725). As in Fig. 1, NO was visualized in the presence of SOD to prevent conversion to peroxynitrite. Protein S-nitrosylation was susceptible to inhibition by L-NPA (Fig. 2C, D \**P =* 0.0152, t = 2.615) but not 1400W, pointing to its dependence on NO generated by NOS1 or NOS3 but not NOS2 (Fig. 2C, D). In contrast, protein nitration was unaffected by L-NPA but was significantly reduced by 1400W, indicating its dependence on NO generated by NOS3, presumably from inflammatory cells, not NOS1 (Fig. 2E,F).

**Figure 2.**
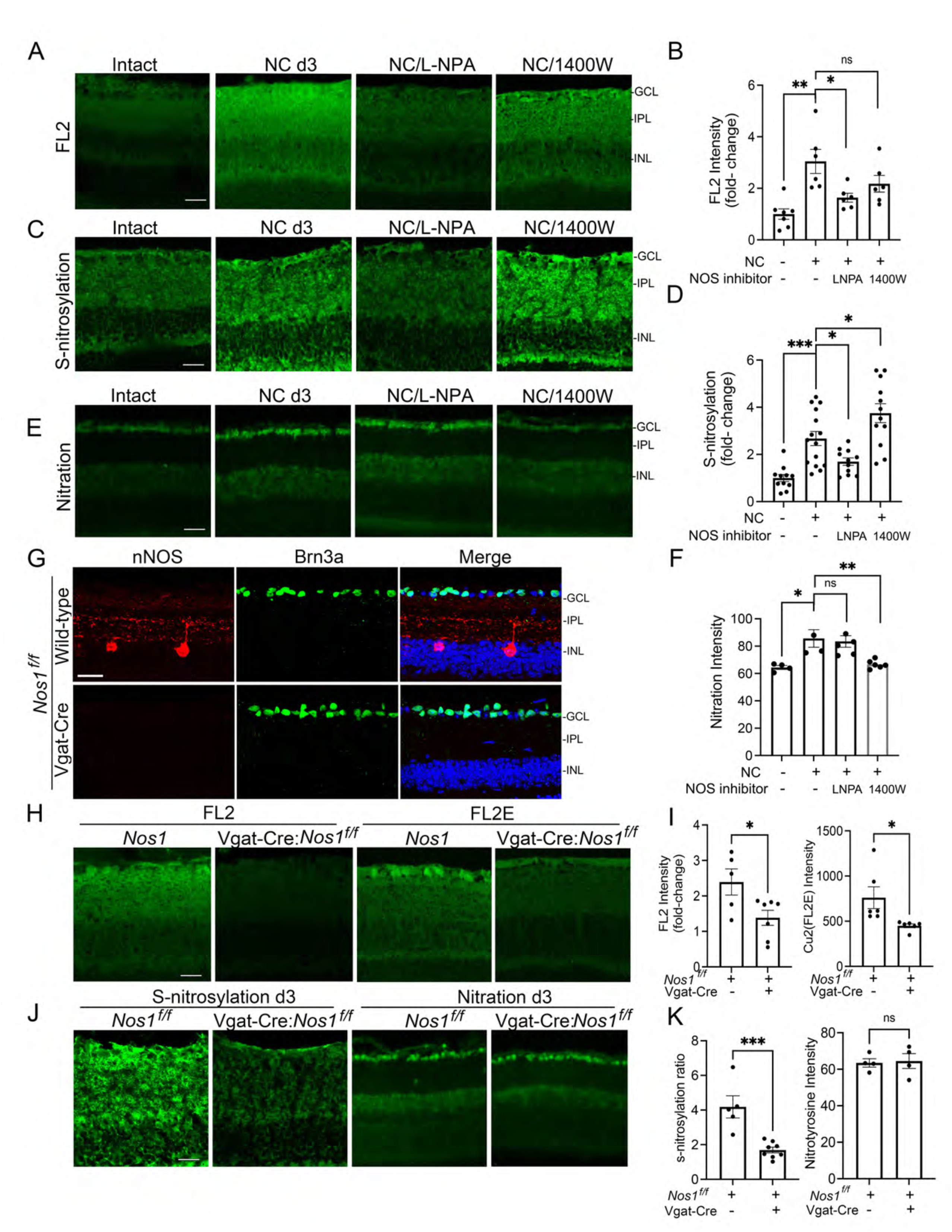
NOS1 as a source of nitric oxide: pharmacological and genetic studies. (A, B) NO elevation detected with FL2 in the retina 3 days after ONC. The signal is strongly sup-pressed by the NOS1/NOS3 inhibitor L-NPA but less so by the NOS2 inhibitor 1400W. Scale bar: 25 µM (A). (B) Quantitation: fold-change normalized to intact control (n = 6-7 mice/ condition; \**P =* 0.0179, t = 2.827, ** *P =* 0.0014, *t* = 4.237; ns = 0.1563, *t* = 1.533). (C, D) Protein S-nitrosylation visualized in the retina 3 days post-ONC. The NOS inhibitor L-NPA prevented protein S-nitrosylation. Quantitation: fold-change normalized to intact control (n = 11-15 mice/condition; \*\*\**P =* 0.0001, t = 4.506; \**P =* 0.0152, t = 2.615 (NC vs. NC+L-NPA); \**P =* 0.0368, *t* = 2.206 (NC vs. NC+1400W). (E, F) Elevation of protein nitration 3 days after ONC. Unlike NO and protein S-nitrosylation, protein nitration is unaffected by L-NPA but is reduced by the NOS2 inhibitor 1400W. Quantitation: fold-change normalized to control samples (n = 4-5 mice/condition; \**P =* 0.0182, *t* = 3.217; \*\**P* < 0.0074, *t* = 3.559; ns = 0.7753, *t* = 0.2953). (G) Ni-tric oxide synthase-1 (NOS1, nNOS: *red*) is expressed in large amacrine cells in the innermost inner nuclear layer (INL) of the retina and in scattered smaller amacrine cells within the INL and ganglion cell layer (GCL, not shown). Amacrine cell-specific deletion of NOS1 (*NOS1^flx/flx^ VGATCre* mice) eliminates the NOS1 signal. Brn3a is a marker for RGCs (*green*). (H, I) Amacrine cell-specific NOS1 deletion reduces the retinal NO signals after ONC as detected by FL2: *left panel*) and FL2E (*Right panel*). Controls show the expected elevation of NO in control Cre^-/-^: NOS1^flx/flx^ mice. Scale bar: 25 µM. Quantitation of results (n = 6-8 mice/condition; \**P =* 0.0294, *t* = 2.538 in left panel; \**P =* 0.0295, *t* = 2.537 in right panel). (J, K) Amacrine cell-specific NOS1 deletion decreases protein S-nitrosylation at D3 after ONC but does not alter protein nitration. Quantitation of results (n = 6-8/condition; \*\*\**P =* 0.0007, t = 4.662 in left panel; n = 4/condition; ns = 0.8377, *t* = 0.2139 in right panel).

As described previously, NOS1 is expressed in subsets of amacrine cells that include small cells within the ganglion cell-and inner nuclear layers (GCL, INL), and large, widely ramifying cells in the INL (44–48). In our hands, NOS1 immunostaining in cross-sections through the normal mouse retina was dominated by the latter cells and what appear to be neuritic processes in the inner plexiform layer (IPL, Fig. 2G, *top panels*). Amacrine cell-specific deletion of NOS1 in *NOS1^flx/flx^ VGAT* mice eliminated cellular NOS1 immunostaining in the INL and IPL (Fig. 2G, *bottom panels)*. As expected, AC-selective NOS1 deletion also suppressed NO elevation in the retina after optic nerve injury, as visualized with both the membrane-impermeable detector FL2 (Fig. 2H, *panels 2 vs. 1* and Fig. 2I, *left*) and the membrane-permeant detector FL2E (Fig. 2H, *panels* 4 vs. 3 and Fig. 2I, *right*). Further, in parallel to the results shown above, AC-selective deletion of NOS1 suppressed protein S-nitrosylation in the retina (Fig. 2J, *panels 2 vs. 1* and Fig. 2K), confirming its dependence on NO generated by amacrine cell NOS1. Protein nitration, on the other hand, was unaffected (Fig. 2J, *panels 4 vs. 3* and Fig. 2K), consistent with the pharmacological results (Figure 2E, F) suggesting that protein nitration following ONC depends on NO generated by NOS3.

### NOS1 inactivation suppresses optic nerve regeneration

We next used pharmacological and genetic approaches to investigate the role of NO in optic nerve regeneration. As shown recently (19), combining RGC-selective deletion of *Pten* with the myeloid cell-derived growth factor oncomodulin (34) and the non-hydrolyzable, membrane-permeable cAMP analog CPT-cAMP induced strong optic nerve regeneration (Fig. 3A, *top*). To test for a possible role for nitric oxide in the regenerative response promoted by *Pten* deletion, CPT-cAMP, pus oncomodulin, we used the NOS1 inhibitor L-NPA at 100 µM, a concentration that suppresses NO elevation. L-NPA diminished the effects of *Pten* deletion/Ocm/CPT-cAMP by ∼ 50% (i.e., halved the difference between regeneration induced by treatment and the negative control (Figs. 3A *bottom* and 3B \*\**P =* 0.0042, t = 3.275). RGC survival, on the other hand, was unaffected (Fig. 3C,D; *NS: P* = 0.9900, t = 0.0128).

**Figure 3.**
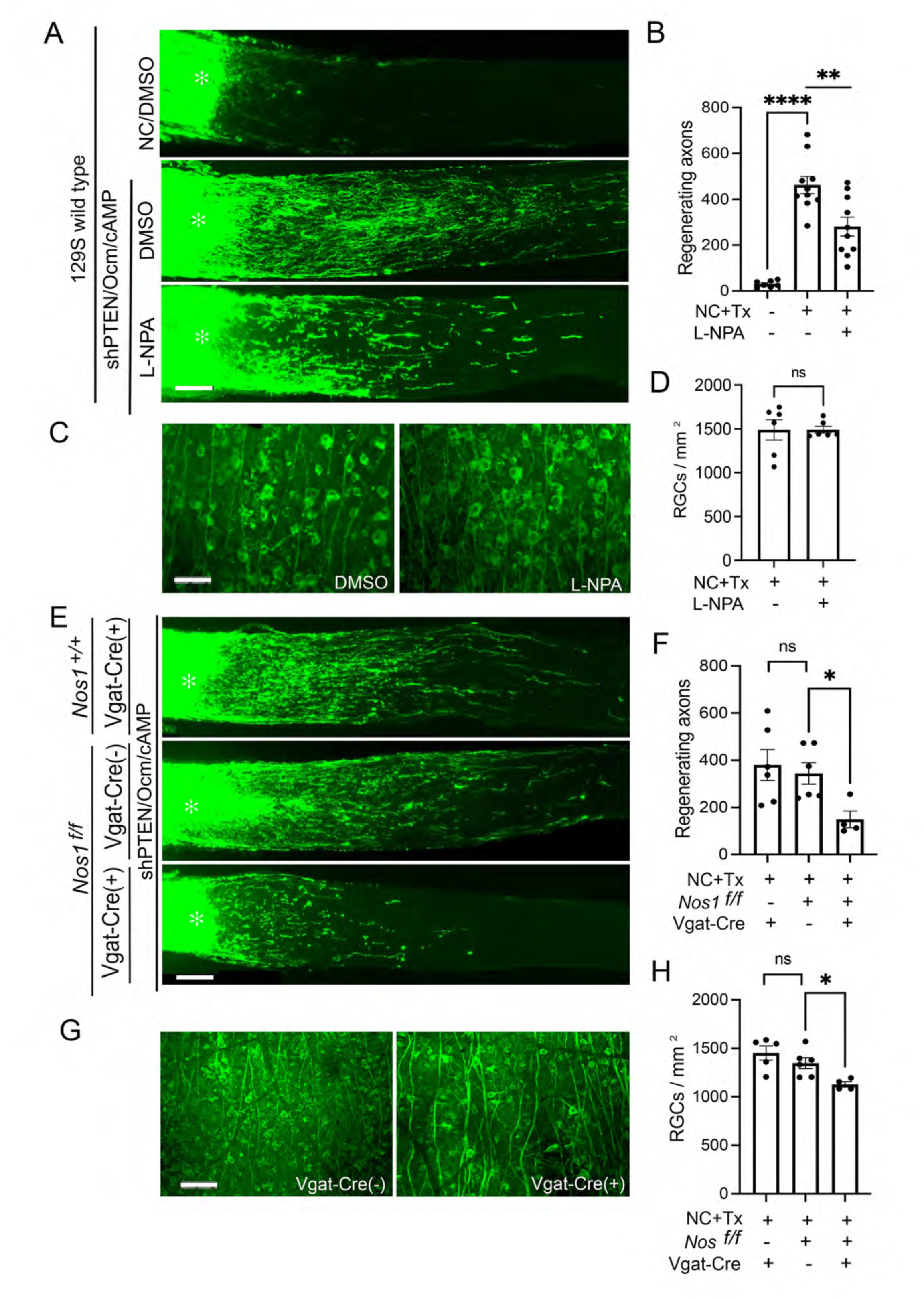
NOS1 Inactivation suppresses optic nerve regeneration. (A) Longitudinal sections through the optic nerves of 129S mice immunostained for the anterograde tracer CTB to visualize axon regeneration after optic nerve injury. Robust regeneration induced by Pten deletion (in-traocular AAV2-shPten virus: shPTEN) combined with oncomodulin (Ocm) and CPT-cAMP (*middle panel*) is suppressed by NOS1 inhibitor L-NPA (*lower* panel). *Asterisks* indicate injury site. Scale bar: 100 µm. (B) Quantitation of regeneration 0.5 mm distal to injury site (n = 7-10 nerves/ condition; \*\**P =* 0.0042, *t* = 3.275; \*\*\*\**P <* 0.0001, *t* = 9.590). (C) Representative areas of retinas immunostained for RGC-selective marker βIII tubulin two weeks after optic nerve injury. Scale bar: 50 µm. (D) Quantification of RGCs survival (n = 6 retinas/condition; ns, *P =* 0.990, *t* = 0.0128). (E, F) Amacrine cell-specific NOS1 deletion reduces optic nerve regeneration. Strong regeneration induced by the combinatorial treatment described in A in two types of littermate control mice (Vgat-Cre^+/-^: NOS1^w-t^ mice, *top panel* and NOS1*^flx/flx^*;Cre^-/-^, *middle* panel) is reduced in mice lacking NOS1 in amacrine cells (NOS1*^flx/flx^*;Vgat-Cre^+/-^ mice, *bottom panel*). Scale bar: 100 µm. Quantitation of regeneration 0.5 mm distal to injury site (n = 4-6 nerves/condition, \**P =* 0.0156, *t* = 3.060; *ns, P =* 0.664, *t* = 0.4474). (G, H) Whereas both types of littermate control mice (*NOS1^flx/flx^ Cre^-/-^* and *NOS1^+/+^ VGAT-Cre*) show high levels of RGCs survival, amacrine cell-specific NOS1 deletion (*NOS1^flx/flx^ VGAT-Cre^+/-^* mice) show a small decrease. Scale bar: 50 µm. Quantitation of results (n = 4-6/condition, \**P =* 0.0176, *t* = 2.979; *(NOS1^flx/flx^ VGAT-Cre^-^*: *vs NOS1^flx/flx^ VGAT-Cre^+/-^*); \*\**P = 0.0073, t = 3.731 (NOS1^wt^ VGAT-Cre^+/-^*: *vs. NOS1^flx/flx^ VGAT-Cre^+/-^*)*; ns, P =* 0.2789, *t* = 1.152).

These last results suggested that NO production by NOS1 after ONC is necessary for the full regenerative response. To test this hypothesis genetically, we examined the consequences of deleting NOS1 in amacrine cells. These experiments employed two controls: mice expressing Cre recombinase and wild-type NOS1 (Fig. 3E, *top panel*); and mice bearing a floxed version (tagged with flanking LoxP-sites) of the NOS1 gene (*NOS1^flx/flx^*) but lacking Cre recombinase (Fig. 3E, *middle panel*). Whereas both controls showed substantial axon regeneration, mice lacking NOS1 in amacrine cells (*NOS1^flx/flx^:VGAT-Cre*, Fig. 3E, *bottom panel*) showed a ∼ 60% decline in regeneration (Fig. 3F \**P =* 0.0156, *t* = 3.060) accompanied by a modest decrease in RGC survival (Fig. 3G,H \**P =* 0.0176, *t* = 2.979).

To test whether diminished regeneration was due specifically to a loss of NO production and to investigate the species of NO that enables regeneration, we carried out rescue experiments, combining amacrine cell-selective NOS1 deletion with either of two distinct NO donors. Nitric oxide exists in three redox forms, NO**^•^**, NO^+^, or NO^−^ (124, 125). Both DETA-NONOate, a donor of NO**^•^,** or Cay10562, a putative NO^+^ donor, administered once shortly after ONC, partially restored regeneration (\**P* = 0.0476, t = 2.229 for DETA-NONOate, \**P* = 0.0126, *t* = 2.858 for Cay10562) (*SI Appendix*, Fig. S1A,B). In addition, both DETA-NONOate and Cay10562 enabled some axon regeneration on their own (*SI Appendix,* Fig. S1B).

The activity of NOS1 and NOS3, but not NOS2, is regulated by intracellular Ca^2+^ interacting with calmodulin (CaM) (126–128). Ca^2+^ can enter neurons via NMDA receptor-associated channels, with which NOS1 is commonly associated in the post-synaptic zone of excitatory synapses (57, 67, 129–132). To investigate whether NMDA receptor activation lies upstream of NO generation, we delivered the highly selective NMDA receptor antagonist D-2-amino-5-phosphopentanoic acid (D-AP5, 100 µM) intraocularly once shortly after optic nerve injury. D-AP5 reduced NO levels in the IPL and GCL by ∼ 60% (as detected with FL2E: Fig. 4A,B, \*\**P =* 0.0034, t = 3.816) and reduced regeneration induced by Pten deletion/Ocm/CPT-cAMP by ∼45% (\**P =* 0.0117, t = 3.150), similar to the effect of L-NPA (Fig. 4C,D).

**Figure 4.**
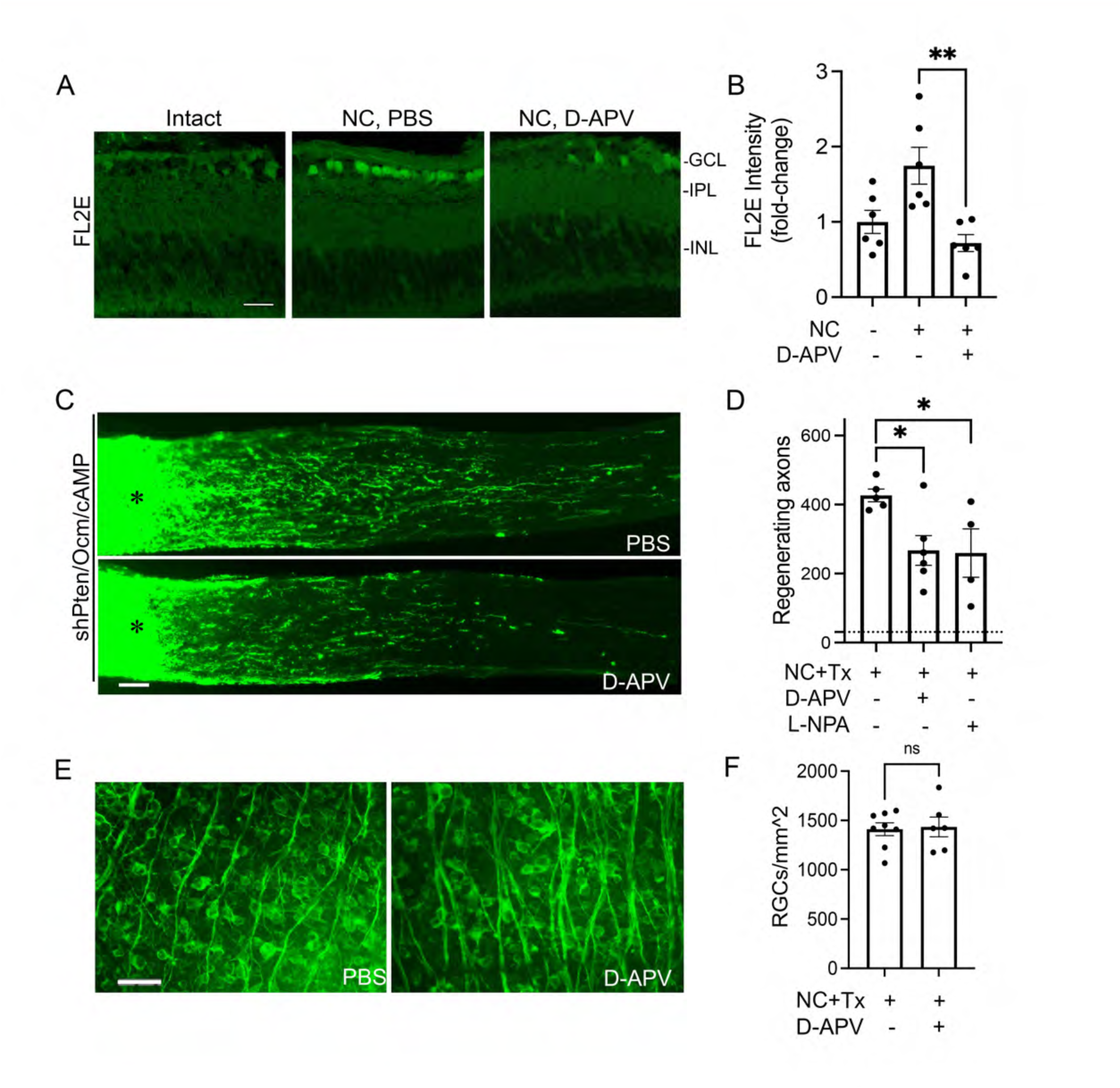
NMDA receptors gate NO elevation and axon regeneration. (A,B) NO elevation after optic nerve crush (NC) in the inner plexiform and ganglion cell layers (IPL, GCL), as visualized with intracellular NO sensor FL2E, is suppressed by NMDA receptor antagonist D-AP5 (1 mM) (A). Scale bar: 25 µm. Quantitation of results (n = 6/condition; \*\**P =* 0.0034, *t* = 3.816). (C, D) Axon regeneration induced by Pten deletion/Ocm/CPT-cAMP is reduced by NMDA-R antagonist D-AP5. Scale bar: 100 µm. Images were taken two weeks after optic nerve crush. *Asterisks* indicate injury site. Quantitation of results (D) 0.5 mm distal to injury site, n = 4-6/condition. \**P =* 0.0117, *t* = 3.150 (*D-AP5 vs. PBS);* \**P =* 0.0371, *t* = 2.568 (*L-NPA vs. PBS*). (E, F) RGCs in representative retinal fields immunostained for βIII tubulin (E). Scale bar: 50 µm. Quantitation of results (F) shows no effect of D-AP5 on RGC survival (n = 6-8/ condition; \*\**P =* 0.8438, *t* = 0.2014).

### The glutamate transporter GLT-1 contributes to the activation of NOS1

The involvement of NMDA receptors in NO elevation after optic nerve injury suggests that extracellular glutamate may increase in the vicinity of amacrine cell NMDA receptors. Extracellular glutamate levels are dynamically regulated by glutamate clearance, mediated primarily by glutamate transporters, (133, 134) and by glutamate release. To define the cellular loci of the glutamate transporters involved in NO generation, we crossed a previously characterized GLT-1 conditional knockout mouse line (82) with either a mouse line expressing GLAST-Cre/ERT (JAX strain #:012586) to obtain Cre- mediated gene deletion in glial cells (135, 136); or with a mouse line expressing VGlut1-Cre to obtain conditional gene deletion in bipolar cells (137). GLT-1 has two major isoforms (84, 138, 139), GLT-1a, which in the retina is reported to be associated with a population of amacrine cells, probably GABAergic (84); and GLT-1b, which is expressed in cone photoreceptors and subpopulations of bipolar cells (84–86). Immunostaining with a high affinity and specific antibody against GLT-1a (82) revealed intense staining above the GCL and in the OPL (Figure 5A, upper panel), as reported previously with an antibody directed against AA 146 to 161 in mouse GLT-1 (EAAT2) (140); in addition there was immunoreactivity in a layer of cell bodies in the INL. These three regions correspond to areas of intense staining using the Mueller cell marker CRALBP. In the corresponding sections from the GLAST-CreERT-driven knockout (Figure 5A, lower panel), the intensity of staining in these three regions was greatly reduced, confirming specificity of the antibody and of GLT-1a localization in Mueller glia.

**Figure 5.**
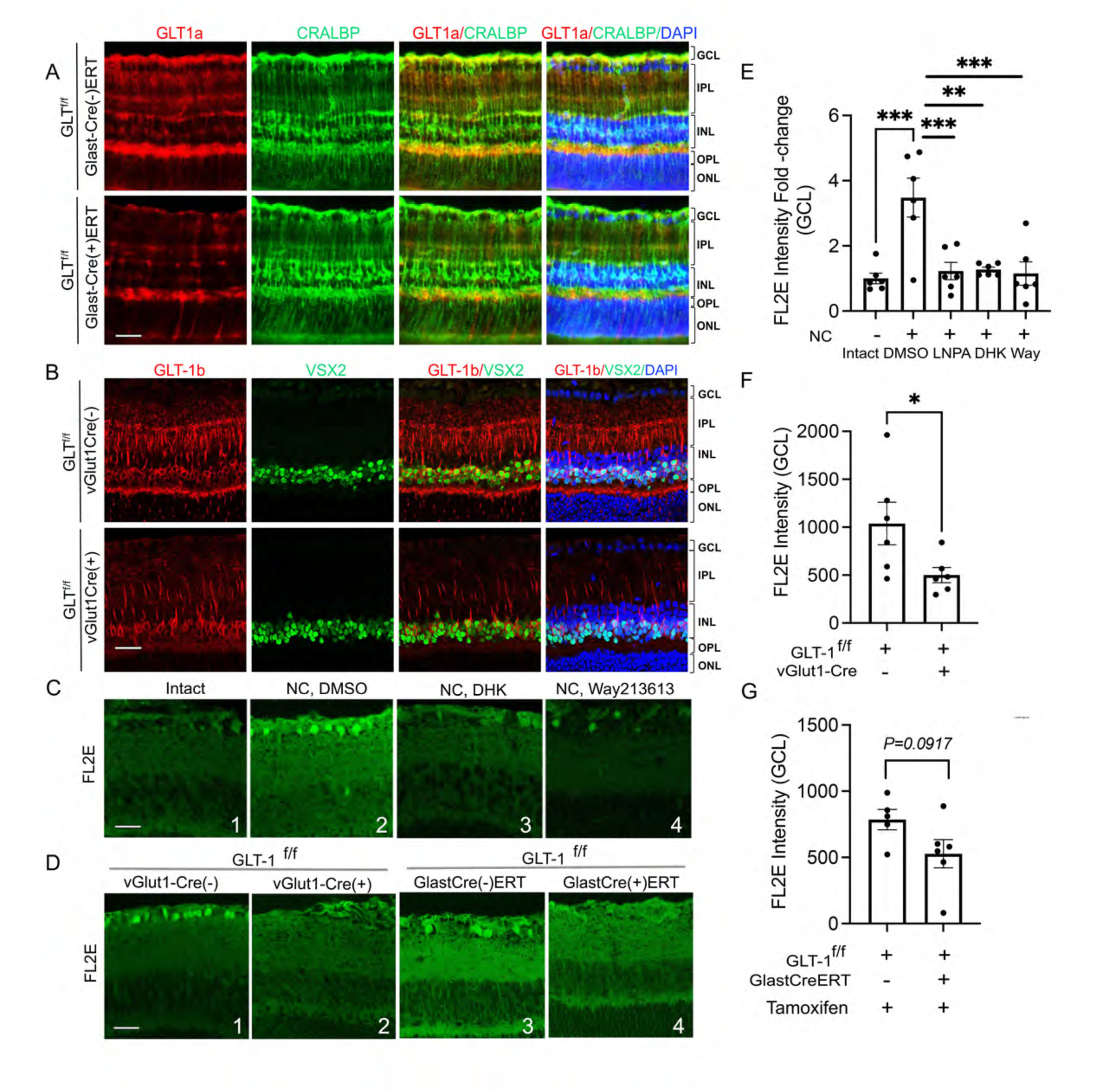
Glutamate homeostasis regulates NO elevation and axon regeneration after optic nerve injury. (A) Immunostaining shows GLT-1a (red) to be expressed in CRALBP-positive Müller glia (green). Crossing GLASTCre-ERT and GLT-1^f/f^ mice suppresses GLT-1a expression following treatment with tamoxifen. (B) Immunostaining shows GLT-1b (red) to be expressed in Vsx2-positive bipolar cells (green). Deletion of GLT-1 in vGlut1Cre;Glt1^f/f^ mice diminishes GLT-1b levels in a subset of bipolar cells. (C) NO elevation (detected with intracellular NO sensor FL2E) is suppressed by glutamate transporter GLT-1 blockers dihydrokainic acid (DHK,1 mM) and WAY 213613 (10 µM) to the same extent as by NOS1 inhibitor L-NPA (not shown). Scale bar: 25 µm. (D) Bipolar cell-specific deletion of GLT-1b in vGlut1-Cre;Glt^f/f^ mice (a, b) but not Müller cell-specific deletion of GLT-1a in GLAST-Cre-ERT;Glt^f/f^ mice (c, d) reduces retinal NO levels (detected by FL2E: a-d; scale bar 25 µm) after ONC. (E) Quantitation of NO levels (visualized with FL2E) in C (n = 6 mice/condition: \*\*\**P =* 0.0002, Intact vs. *ONC/DMSO*; \*\*\**P =* 0.0008, *NC/DMSO vs. NC/L-NPA*; \*\**P =* 0.0010, *NC/DMSO vs. NC/DHK;* \*\*\**P =* 0.0005, *NC/DMSO vs. NC/ Way 213613*)(one-way ANOVA with Tukey’s multiple comparison test; F (4, 25) = 9.259)]. (F) Quantitation of NO levels in GCL in Da, b (n = 6 mice/condition; \**P =* 0.0461, *t* = 2.277, unpaired *t*-test. (G) Quantitation of NO levels (visualized with FL2E) in Dc, d (n = 6 mice/condition; ns, *P =* 0.0917, *t* = 1.887, unpaired t-test).

Similar methods verified the expression of GLT-1b in bipolar cells. Thus, a specific antibody against GLT-1b (141) (Figure 5B, upper panel) revealed intense labeling in radial fibers extending part-way into the IPL, in cells in the outer half of the INL that partly overlap with the bipolar cell-selective marker Vsx2 (142, 143), and in the outer plexiform layer (OPL). Bipolar cell-selective depletion of GLT-1 (*GLT-1^flx/flx^ VGlut1-Cre* mice; Fig. 5B, lower panel) eliminated most of the staining in the IPL and OPL but not in the INL, confirming that the GLT-1b-positive processes in the neuropil are associated with bipolar cells.

### Bipolar cell GLT-1b contributes to NO elevation and optic nerve regeneration

To test the possible role of GLT-1 in altering extracellular glutamate and NMDA receptor activation following ONC (Figure 5C,E), we carried out intraocular injections of either dihydrokainate (DHK) (Ki 23 µM for GLT-1 (EAAT2) and >3 mM for GLAST (EAAT1), EAAC1(EAAT3) or WAY-213613 (IC50 values are 85, 3787 and 5004 nM for EAAT2, EAAT3 and EAAT1, respectively) (144–146), two relatively selective GLT-1 inhibitors. Each GLT-1 inhibitor reduced NO levels as effectively as the NOS1 inhibitor L-NPA [Fig. 5C,E \*\*\**P =* 0.0002, Intact vs. ONC/DMSO; \*\*\**P =* 0.0008, ONC/DMSO vs. ONC/L-NPA; \*\**P =* 0.0010, ONC/DMSO vs. ONC/DHK; \*\*\**P =* 0.0005, ONC/DMSO vs. ONC/ Way 213613)(one-way ANOVA; F (4, 25) = 9.259)]. In complementary genetic studies, bipolar cell-selective depletion of GLT-1 in *VGlut1-Cre*:*GLT-1^flx/flx^*mice suppressed the elevation of NO that would otherwise occur within an hour after optic nerve injury (Fig. 5D-1,2 & F; \**P =* 0.0461, t = 2.277). GLAST-CreERT driven knockout of GLT-1 in Mueller cells (Figure 5D-3,4 & G) produced a trend towards decreased NO production that did not attain statistical significance (Figure 5G). Taken together with the pharmacological studies, these results establish that activation of NOS1 in the retina following optic nerve injury requires GLT-1, and that the isoform GLT-1b, highly expressed in bipolar cells, plays an important, perhaps dominant, role in the glutamate homeostasis dysregulation that leads to NO generation.

Because dysregulation of GLT-1-mediated glutamate transport lies upstream of NO synthesis, we would predict that this dysregulation contributes to enabling robust optic nerve regeneration, a hypothesis that we tested both pharmacologically and genetically. WAY213613 (10 µM, the concentration found to reduce NO levels to baseline: Fig. 5C,E), suppressed regeneration induced by Pten deletion/Ocm/CPT-cAMP by ∼ 50% (Fig. 6A-1,3; 6B \*\**P =* 0.0034, t = 3.396) and diminished RGC survival by ∼ 40% (Fig. 6A-2,4; 6C \*\**P =* 0.0089, t = 3.589). In genetically engineered mice, GLT-1 deletion in bipolar cells (*VGlut1-Cre*:*GLT-1^flx/flx^*) showed a 37% decrease in regeneration compared to two types of littermate controls, *i.e*., mice expressing Cre recombinase in bipolar cells on a wild-type GLT-1 background (*GLT-1*^wt/wt^ *VGluT1-Cre*); and mice expressing *GLT-1* with loxP elements but not Cre recombinase (*GLT-1*^flx/flx^) (Fig. 6D-1,3 and 6E \**P =* 0.0493, t = 2.135). There was no significant difference between the two control populations, which were therefore pooled for purposes of statistical analysis. Bipolar cell-specific deletion of GLT-1 also resulted in a 25% decline in RGC survival (Fig. 6D-2,4; 6F: \*\*\**P =* 0.0005, t = 4.235). GLT-1 deletion in Mueller glia (*GLAST-Cre*:*GLT-1^flx/flx^* mice: Fig. 6G,H) resulted in a 42% decrease in regeneration compared to control mice (*GLT-1^fl/fl^* mice not expressing GLAST-CreERT but treated with tamoxifen: \*\*\*\**P* <0.0001, *t* = 8.874, unpaired t-test).

**Figure 6.**
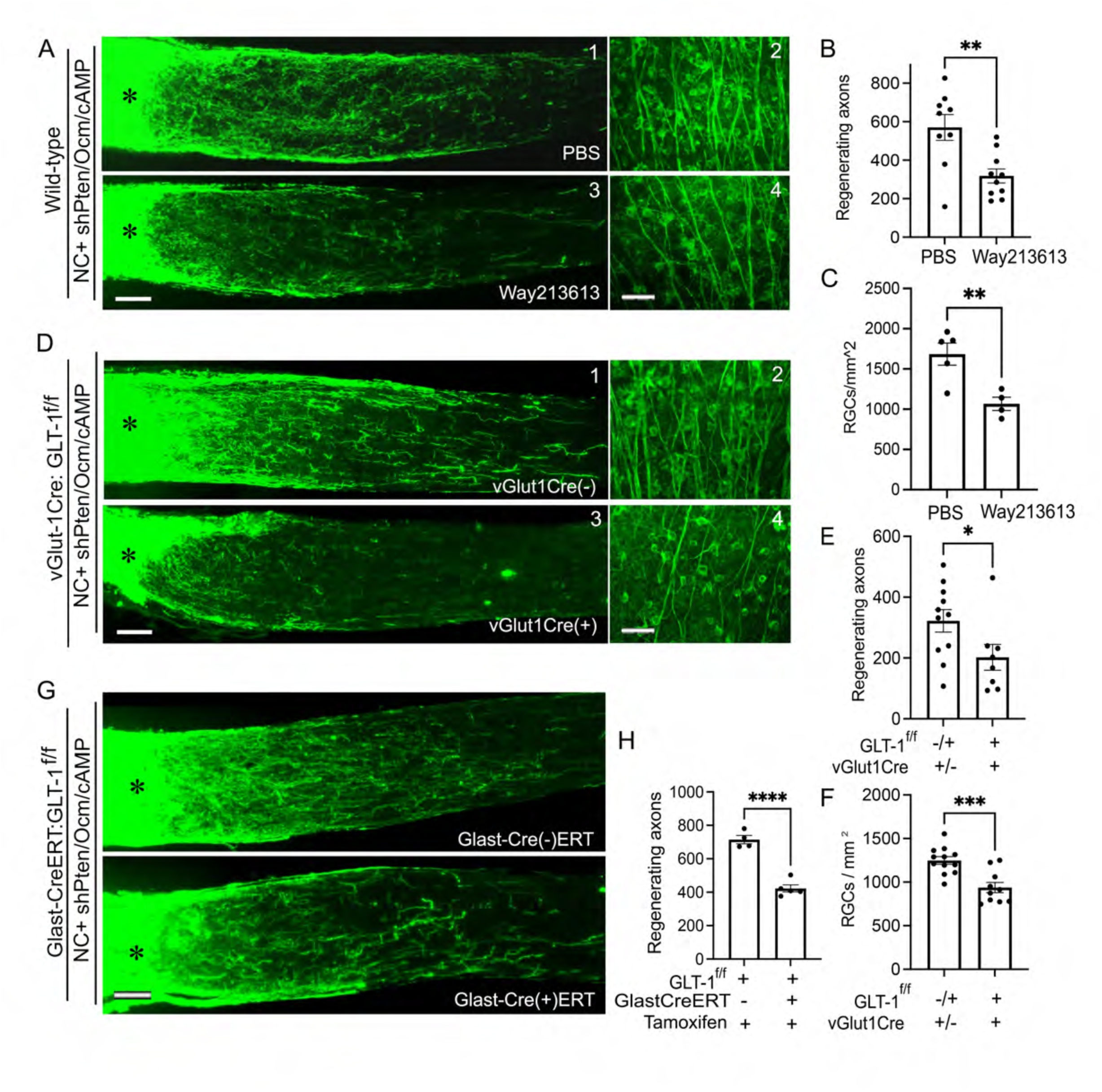
Glutamate homeostasis gates optic nerve regeneration. (A) Axon regeneration in-duced by *Pten* deletion/rOcm/CPT-cAMP in 129S wild-type mice 2 weeks after optic nerve injury (*left panel*, a) is diminished by GLT-1 inhibitor WAY213613 (left, *lower panel,* c). (Scale bar: 100 µm. *Asterisks* indicate injury site). Representative retinal fields (b,d) show RGC survival after treatment without (*top,* b) and with (*lower,* d) GLT-1 inhibitor Way213613 (Scale bar: 50µm.) (B) Quantitation of axon regeneration 0.5 mm distal to injury site (n = 9-10/condition; \*\**P =* 0.0034, *t* = 3.396). (C) Quantitation of RGC survival in whole-mount retina as described in A (n = 4-5/condition; \*\**P =* 0.0089, *t* = 3.589). D) Bipolar cell-specific deletion of GLT-1b in vGlut1-Cre:Glt^f/f^ mice reduces axon regeneration induced by *Pten* deletion/rOcm/CPT-cAMP after ONC (left panel a, c. scale bar: 100 µm *Asterisks* indicate injury site) and RGC survival after ONC (right panel b, d. Scale bar: 50 µm). (E) Quantitation of axon regeneration as described in D at 0.5 mm distal to injury site. (n = 8-11/condition; \**P =* 0.0495, *t* = 2.135, unpaired *t*-test). (F) Quan-titation of RGC survival as described in D (n = 10-12/condition; \*\*\**P =* 0.0005, *t* = 4.235). (G) Müller glia-specific deletion of GLT-1 in GlastCreERT:Glt^f/f^ mice after 5 days of tamoxifen treat-ment diminishes axon regeneration (scale bar: 100 µm. *Asterisks* indicate injury site). (H) Quanti-tation of axon regeneration in G (n = 5/condition, \*\*\*\**P* <0.0001, *t* = 8.874, unpaired t-test).

Because NO activates guanylate cyclase (GC), the enzyme that generates the second messenger cyclic GMP (cGMP), we investigated whether the NO-GC-cGMP pathway contributes to robust optic nerve regeneration. Using an antibody against cGMP, we found increased labeling of cells in the GCL following ONC at 1 day and 3 days (compared with control retinas), which was blocked by injection of L-NPA (Figure 7A,C). However, inactivation of NOS1 in amacrine cells did not alter cGMP labeling in the GCL (Figure 7B,D). In addition, neither the specific NO-dependent GC inhibitor 1H-[1,2,4]oxadiazolo[4,3-a]quinoxalin-1-one (ODQ) or the protein kinase G inhibitor Rp-cGMP altered regeneration induced by Pten deletion/Ocm/CPT-cAMP (Fig. 7E,F).

**Figure 7.**
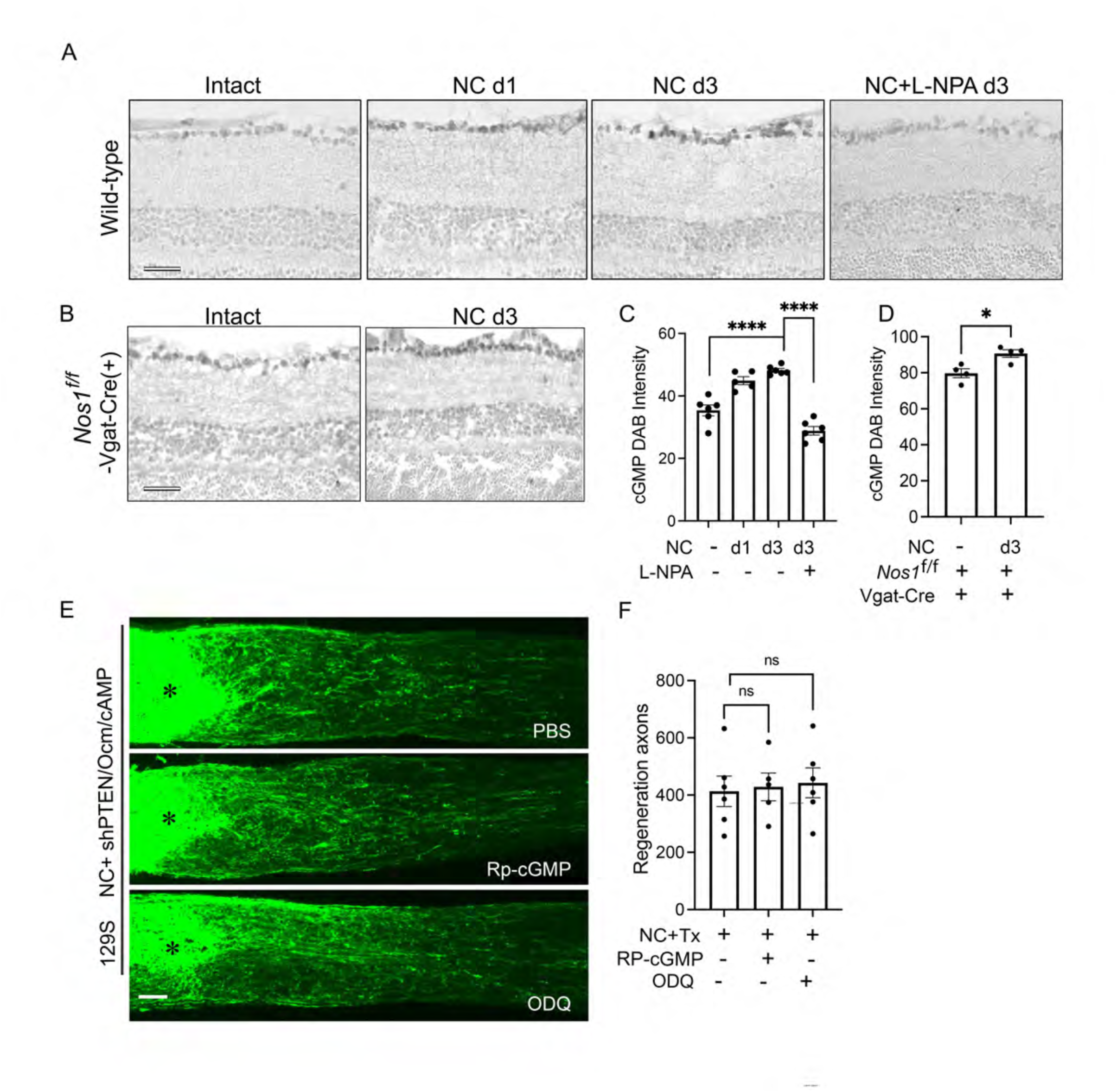
NO does not regulate regeneration through guanylate cyclase. (A) GMP content, as evaluated by DAB immunocytochemistry, increased continuously in RGC layer after optic nerve injury but was blocked by NOS1 inhibitor L-NPA (applied to wild type mice prior to optic nerve crush). (B) Amacrine cell-selective NOS1 deletion (NOS1*^flx/flx^*:Vgat-Cre) did not diminish nerve crush-induced cGMP elevation. (C, D) Quantitation of cGMP DAB results for conditions shown in A and B: n = 6/condition; \*\*\*\**P <* 0.0001, *t* = 7.044 (*Intact vs NC d3*); \*\*\*\**P =* 0.0001, *t* = 13.25 (*NC d3 vs NC/LNPA d3*) in C; n = 4/condition; \**P =* 0.0149, *t* = 3.377 in D). (E) Axon regeneration induced by Pten deletion combined with rOcm and CPT-cAMP was unaltered by protein kinase G inhibitor Rp-cGMP or by guanylate cyclase inhibitor ODQ. Scale bar: 100 µm. *Asterisk* indicates injury site. (F) Quantitation of regeneration 0.5 mm distal to injury site. n = 5-6 condition: ns, *P =* 0.8367, *t* = 0.2121 (PBS *vs*. RP-cGMP); *P =* 0.6982, *t* = 0.3992 (PBS *vs.* ODQ).

Going further upstream in the NO generation pathway, we considered phenomena that might alter glutamate transport in bipolar cells. The direction (and magnitude) of glutamate flux through glutamate transporters is thermodynamically coupled to the electrochemical gradients of Na^+^, K^+^, and protons across the plasma membrane that power the work of concentrating glutamate (69, 147, 148). Because optic nerve injury alters expression of several K^+^ channels (149), and because damage to other types of neurons is known to cause p38-dependent phosphorylation, translocation, and activation of the voltage-gated K^+^ channel Kv2.1 (100–102), we examined the possible involvement of Kv2.1 in NO elevation. We observed a modest but significant increase in Kv2.1 phosphorylation as early as 1 hr after ONC (Fig. 8A,B, \*\*\**P =* 0.001, t = 4.618), and two Kv2.1 blockers, RY796 (10 μM) and stromatoxin (5 μM) both suppressed NO elevation after optic nerve injury [Fig. 8C-E, *P = 0.0495, t = 2.235 (NC vs. RY796.); **P = 0.0043, t = 3.675 (NC vs stromatoxin)].

**Figure 8.**
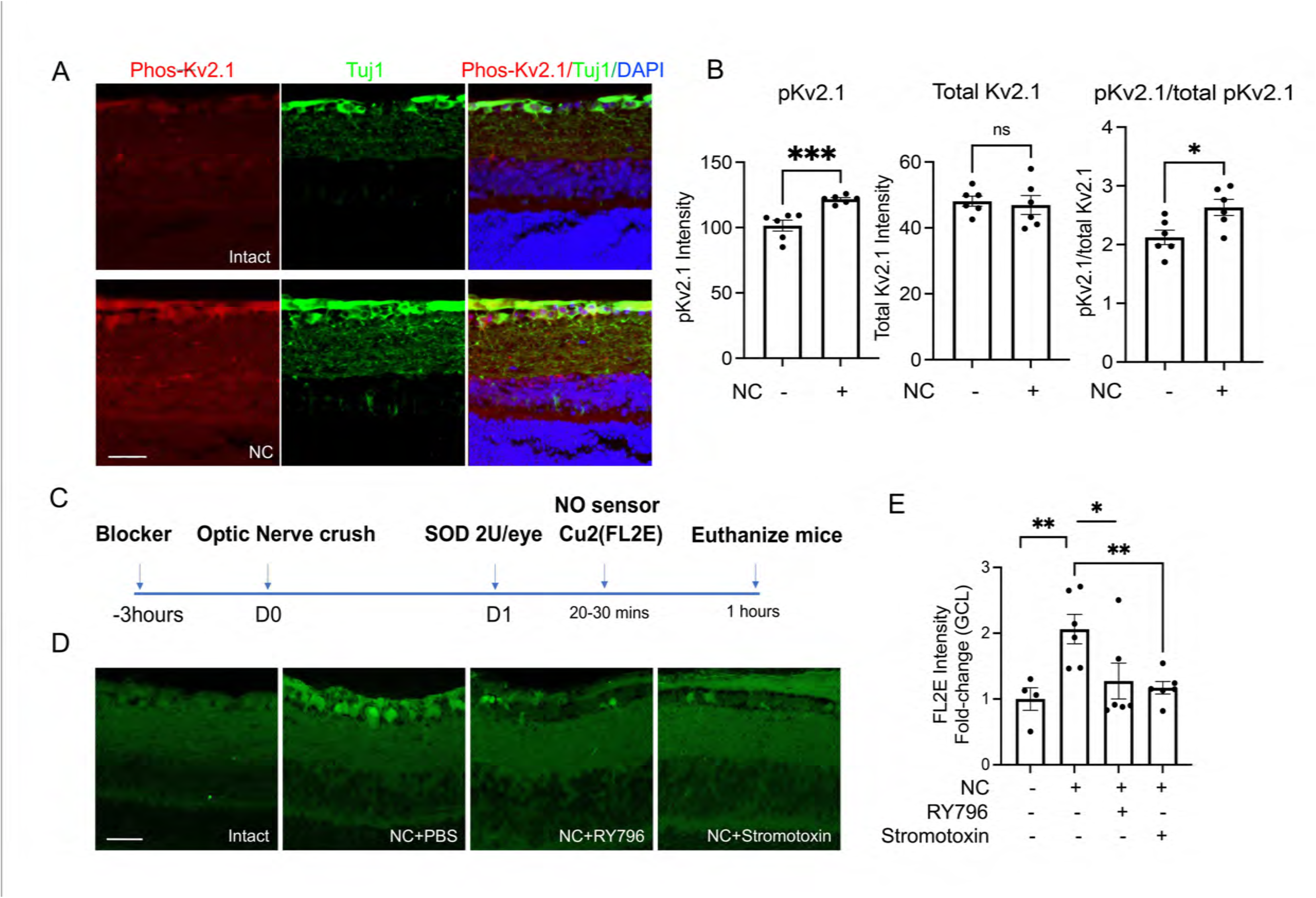
Voltage-regulated K^+^ channel Kv2.1 regulates retinal NO levels. (A, B) Immunostaining (A) shows elevation of p38-mediated phosphorylated K^+^ channel Kv2.1 (pKv2.1, *red*) in RGCs (counterstained with antibody Tuj1 against βIII tubulin, *green*) after optic nerve injury. Scale bar: 25 µM. Quantitation of changes (B). n = 6/condition; \*\*\**P =* 0.001, *t* = 4.618; ns, *P =* 0.7302, *t* = 0.3546; \**P =* 0.0202, *t* = 2.757). (C) Timeline of NO signal detected by FL2E. DMSO control or Kv2.1 channel blockers RY796 or stromatoxin were administered 3 h before optic nerve crush (ONC). SOD1 was administered the following day, followed 20-30 min later with the NO sensor FL2E. Mice were euthanized 1 h later and processed to visualize NO signal. (D, E) NO signal (FL2E) is elevated in GCL after optic nerve crush and is blocked by both RY796 and stromatoxin. Scale bar: 25µM. Quantitation of NO signal in GCL (E), n = 4-6/condition, \*\**P =* 0.009, *t* = 3.427 (NC vs Intact); \*\**P =* 0.0043, *t* = 3.675 (NC with *vs*. without stromatoxin); \**P =* 0.0495, *t* = 2.235 (NC with *vs*. without RY796).

Finally, we previously showed that optic nerve injury results in an increase in an autometallographic signal (AMG) in the IPL of the retina that is suppressed by Zn^2+^ chelators; and that these chelators improve RGC survival and promote varying degrees of axon regeneration (42). Because NO or an NO derivative can liberate Zn^2+^ from metallothionein by S-nitrosylation (150–153), we investigated whether NO generation lies upstream of Zn^2+^ elevation. In agreement with our previous findings (42), autometallography revealed an increase in signal intensity, suggesting Zn^2+^ elevation in the inner retina after optic nerve injury in wild-type mice (Fig. 9A). Intraocular injection of DETA-NONOate also produced an elevation in the AMG signal, confirming its dependence on NO signaling (Fig. 9A). The elevation in AMG intensity following ONC was diminished by the NOS1 inhibitor L-NPA (10 µM: (Fig. 9A,C \**P =* 0.015, t = 3.086) as well as in mice lacking NOS1 in amacrine cells (*Vgat-Cre* mice: *NOS1^flx/flx^* Fig. 9D,E, \*\*\**P =* 0.0008, *t* = 9.059). Thus, although NO elevation enables the pro-regenerative treatment used here to promote strong axon regeneration, NO elevation also has a potentially deleterious effect in stimulating mobile Zn^2+^ elevation. We queried whether the regeneration observed with Zn^2+^ chelation might itself be dependent upon the release of NO. Combining Zn^2+^ chelation with L-NPA resulted in the same level of regeneration as Zn^2+^ chelation alone, suggesting that the regeneration observed following Zn^2+^ chelation is not dependent upon NOS1 activation.

**Figure 9.**
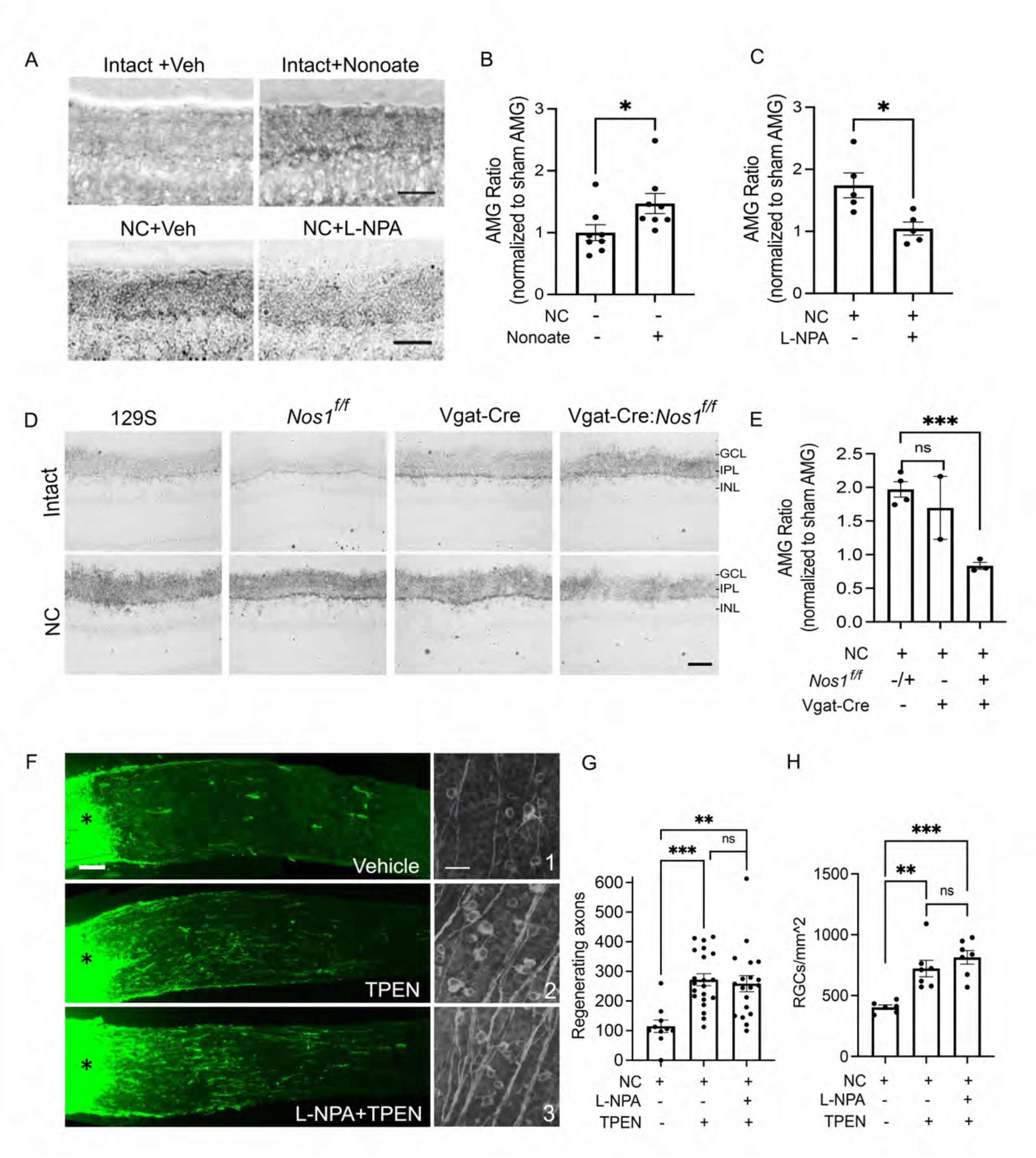
Nitric oxide increases retinal autometallographic (AMG) signal after optic nerve crush and NOS1 inhibitor L-NPA did not alter the benefits of zinc chelation. (A-C) The intensity of the AMG signal is elevated in the inner plexiform layer (IPL) within 1 day after optic nerve crush in wild-type mice. Intraocular injection of NO donor DETA-NONOate (10 μM) without nerve crush increases the AMG signal similar to the effect of optic nerve injury. AMG signal elevation after optic nerve crush is reduced by NOS1 inhibitor L-NPA. Quantification of AMG signal normalized to the signal in sham-operated mice (C). n = 5-8/condition, \**P =* 0.0396, *t* = 2.269 (*Nonoate vs. intact*); \**P =* 0.015, *t* = 3.086 (*NC+PBS vs. NC+L-NPA*). (D, E) AMG signal normalized to the contralateral eye is elevated after unilateral optic nerve crush in 129s wild-type mice and in two types of littermate controls (NOS1^flx/flx^;Cre^-/-^ and NOS1*^wt/wt^*;Vgat-Cre mice), but decreases in NOS1 conditional knockout mice (NOS1*^flx/flx^*;Vgat-Cre^+^). Quantitation of the AMG signal in the IPL (E). n = 2-4/condition; ns, *P =* 0.6611, *t* =0.5709 (NOS1*^wt/wt^* or *NOS1^flx/flx^;Vgat-Cre^-^ vs. NOS1^wt/wt^;Vgat-Cre+*); \*\*\**P =* 0.0008, *t* = 9.059 (*NOS1^flx/flx^ Vgat-Cre^-^ vs. NOS1^flx/flx^;Vgat-Cre^+^*). (F). TPEN, a chelator of Zn^2+^ (and possibly other divalent cations other than Ca^2+^) increased axon regeneration after optic nerve injury, and this effect was not altered by the NOS1/3 inhibitor L-NPA (G). Quantitation of regeneration: n = 19-21/condition, ns, *P* = 0.9060, *t* = 0.3948 (*NC+TPEN vs NC+L-NPA/TPEN*) (H) Quantitation of RGC survival for conditions shown in F. L-NPA alone slightly improved RGC survival after optic nerve injury (n = 5-6/condition, \*\**P* = 0.0079, *t* = 3.511 (*NC+L-NPA vs. NC+PBS*) but did not further enhance RGC survival effect of TPEN. (n=6/condition, ns, *P* = 0.3967, t = 0.8854 (*NC+TPEN vs NC+L-NPA/TPEN*).

## Discussion

### A multicellular signaling pathway generating NO regulates the regenerative response of RGCs to axonal injury

Our results show that optic nerve injury leads to a cascade of events that involves several retinal cell types and that enables RGCs to mount a vigorous regenerative response. Levels of NO increase in the retina within an hour after optic nerve injury and remain high for several days. Of the multiple NO synthase isoforms and cell types that can generate NO, our studies point to NOS1-positive amacrine cells as being the principal source. Amacrine cell-specific deletion of NOS1 reduced retinal NO levels close to baseline levels after optic nerve injury and suppressed the effects of a strongly pro-regenerative treatment, as did the NOS1 inhibitor L-NPA. Upstream events regulating NOS1 activity include activation of NMDA-type glutamate receptors driven by reversal of GLT-1-mediated glutamate transport in bipolar and glial cells. Accordingly, both NO levels and axon regeneration are inhibited by the NMDA receptor antagonist D-AP5, the GLT-1 inhibitors DHK and WAY213613, and by deletion of GLT-1 in bipolar and glial cells. Fig. 10 schematically illustrates the multicellular signaling cascade that is initiated by optic nerve injury and gates RGCs’ ability to regenerate injured axons via NO generation.

**Figure 10.**
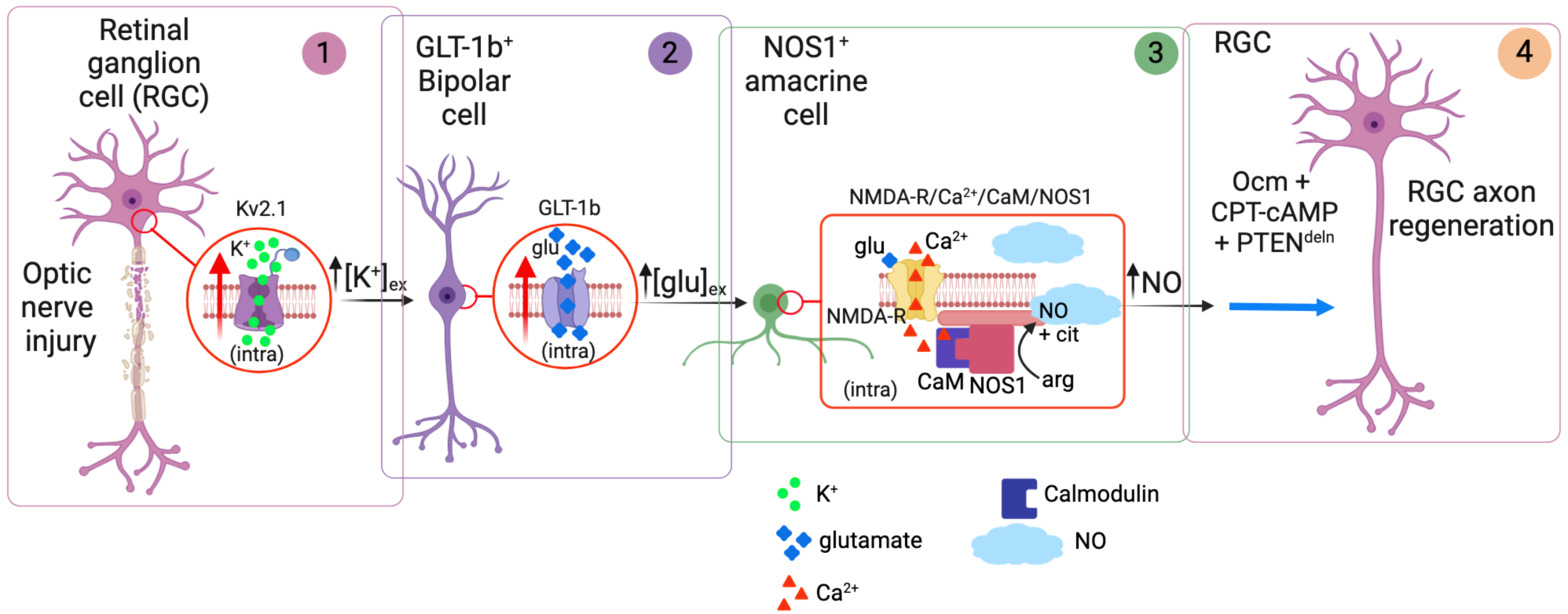
NO pathways gate axon regeneration after optic nerve injury. Retinal ganglion cells (RGCs) cannot normally regenerate axons after optic nerve injury and soon begin to die due to both cell-autonomous and non-cell autonomous factors. (1) Loss of intracellular K^+^ through the voltage-gated channel K_v_2.1 may lead to extracellular ionic imbalance. (2) Glutamate transporter GLT1b in bipolar cells normally takes up extracellular glutamate to maintain low levels of the transmitter, thus enabling precise intercellular signaling. However, transport is blocked or reversed after optic nerve injury, likely due to changes in extracellular ion balance. (3) Elevation of extracellular glutamate activates NMDA receptors, allowing an influx of Ca^2+^ that interacts with intracellular calmodulin to stimulate NOS1 and generate NO. (4) NO gates axon regeneration induced by combining PTEN deletion, rOcm and cPT-cAMP.

### Prior studies of involvement of nitric oxide in response to injury in the eye

Our results differ from prior studies in several ways. In rats, NOS1-derived NO has been reported to promote RGC death (50), whereas in goldfish, studies using retinal explants reported modest neurite outgrowth in response to high concentrations of an NO donor, with RGCs themselves being the physiological source of NO (53). In Drosophila, elevated NO leads to axon pruning and suppresses axon regeneration during development (52). In contrast to these reports, our results highlight the intercellular signals regulating the generation of nitric oxide after injury and point to AC-derived NO as a positive regulator of mouse optic nerve regeneration.

### NOS1 expression in a subpopulation of amacrine cells in the retina, and its regulation by NMDA receptor activation after ONI

Within the retina, NOS1 is expressed in a sparse population of large diameter amacrine cells (ACs) in the inner nuclear layer (INL) having wide ramifications in the inner plexiform layer (IPL); and in sparse populations of smaller ACs in the inner nuclear and ganglion cell layers (44–48, 154). NOS1 exists in 3 isoforms, of which NOS1α dominates in the retina (46). Endothelial NOS (NOS3) expression is restricted to blood vessels and the outer limiting membrane, whereas inducible NOS (NOS2) is largely undetectable in the normal retina (46, 48). NOS1-positive ACs express AMPA/kainate-and NMDA-type glutamate receptors, and normally modulate NO levels and neural activity in response to changes in ambient light (48, 154–156). Ca^2+^ entry through NMDA receptor channels enables formation of a Ca^2+^-calmodulin complex that activates NOS1. Accordingly, the NMDA receptor antagonist D-AP5 was found to block NO production and, as with NOS1 deletion, to suppress axon regeneration without affecting RGC survival.

### Downstream signaling pathways mediating effects of nitric oxide on regeneration

Nitric oxide is a highly reactive free radical gas, the production of which has multiple consequences for cellular physiology (157–159). In our study, visualization of NO *per se* required exogenous superoxide dismutase, an enzyme that converts superoxide to O_2_ and H_2_O_2_ (160).

Optic nerve injury induces superoxide production in RGCs (161, 162), which represents a likely basis for the rapid conversion of NO to peroxynitrite implied by our results (163). Increased peroxynitrite formation aligns with the increase in protein nitration that we observe for at least a week after optic nerve injury (164, 165), although alternate mechanisms downstream of nitric oxide and superoxide production are possible (166). In addition to protein nitration, we also observed elevated protein S-nitrosylation following optic nerve injury. S-nitrosylation was eliminated following treatment with L-NPA or in animals in which NOS1 was specifically inactivated in amacrine cells, indicating its dependence on NO generation by NOS1 and raising the possibility that S-nitrosylation of one or more proteins underlies the observed effect of NO production by NOS1 on regeneration. One protein that is inactivated by S-nitrosylation is PTEN, which might be expected to enhance axon regeneration (4, 167). However, in the present studies, PTEN is already deleted in RGCs as a component of our pro-regenerative therapy, and thus PTEN S-nitrosylation is unlikely to contribute further to regeneration. Nonetheless, it remains possible that S-nitrosylation of other proteins, including other components of the mTOR pathway (168), could be important in this regard. In contrast to S-nitrosylation, protein nitration was inhibited by the NOS2 inhibitor 1400W, but was unaffected by L-NPA and NOS1 deletion in ACs, implying its dependence upon NO production by NOS2, an enzyme found in inflammatory cells that can produce very high levels of NO (169, 170). It is noteworthy that, despite the production of nitric oxide by NOS2, nitric oxide production by NOS1 is required to promote regeneration following ONI. This apparent paradox suggests that highly localized NO signaling is likely to be critical for promoting both the protein S-nitrosylation and regenerative responses that are observed (171).

Based on several studies linking cGMP to regeneration, another signaling pathway that might be hypothesized to mediate the pro-regenerative effects of NO elevation is activation of guanylate cyclase, production of cGMP, and stimulation of cGMP-dependent protein kinase G (53, 172). However, the absence of an effect of inhibiting guanylate cyclase, the NO-activated enzyme that generates cGMP, or of the PKG antagonist Rp-cGMP, makes this possibility unlikely.

### The glutamate transporter GLT-1 regulates NMDA receptor activation and NO production

Because the EC_50_ of the NMDA receptor for glutamate is ∼2 μM (72, 173), and because activation of NMDA receptors can profoundly affect neural activity and pathology, the ambient glutamate concentration in the vicinity of NMDA receptors is expected to be much lower than 2 µM, and, in the hippocampus, is estimated to be ∼25 nM (55). The elevation in extracellular glutamate required to activate NMDA receptors, as evidenced by the role of these receptors in NO generation, could arise through increased synaptic release or through a perturbation of glutamate transport--either decreased clearance of extracellular glutamate or reversal of transport (64–69), among other possibilities including release from glutamate-permeable, volume-sensitive channels in astrocytes (174–178). Like other glutamate transporters, GLT-1 takes up glutamate to maintain low levels in the extracellular space (55, 71). Unlike GLT-1a expression in brain neurons, which appears to be expressed only in excitatory axon terminals (80–82), GLT-1b is expressed in bipolar cells in a somatodendritic pattern (Figure 5B). GLT-1b, but not GLT-1a, has a PDZ protein interaction domain at its C-terminus that allows it to interact with synaptic proteins such as PICK1 (179, 180) and which may be important in regulating the expression, trafficking, and function (181) of GLT-1 at the plasma membrane of bipolar cells. Since glutamate transporters can either be a source of a sink for extracellular glutamate, blockers of glutamate transporters [for definition of “blocker”, see section 6.5 in ref. (134)] may either raise or lower extracellular glutamate, depending on the circumstances. We found that either of two GLT-1 blockers reduced NO elevation after optic nerve injury and suppressed axon regeneration to a similar extent as deleting NOS1 in amacrine cells or intraocular administration of D-AP5; genetic deletion of GLT-1 in bipolar cells had a nearly similar effect. These results imply that reversal of uptake of glutamate by GLT-1b contributes to events leading to NO generation and the gating of optic nerve regeneration. Deleting GLT-1 in Mueller cells also decreased axon regeneration, consistent with the GLT-1 expression pattern found in recent histological and transcriptomic studies (140, 182, 183).

There has long been interest in the possible role of changes in extracellular K^+^ in modulating glutamate clearance, especially in the setting of ischemia (184–186), with recent studies showing an inhibition of glutamate clearance by K^+^ released as a consequence of neuronal activity (91–93). RGCs undergo apoptosis after optic nerve injury (187), and, in other types of neurons (100–102), the loss of intracellular K^+^ following phosphorylation-dependent Kv2.1 translocation to the plasma membrane has been shown to be a critical step in the apoptotic pathway. Our results point to a modest but rapid phosphorylation of this channel in RGCs within an hour after optic nerve injury, and show that two different Kv2.1 inhibitors, RY796 and stromatoxin, inhibit NO elevation post-injury. These results suggest, then, that Kv2.1 channels may be the source of extracellular K^+^ driving glutamate release mediated by GLT-1 expressed in bipolar cells and in Mueller cells following optic nerve damage. Further work will be required to characterize the signaling pathway(s) underlying Kv2.1 phosphorylation in RGCs after axonal injury (102).

Nearly thirty years ago, Garthwaite and colleagues showed in mixed rat neuron-glial striatal cultures that, following a brief excitotoxic insult, dysregulation of glutamate homeostasis, presumed to be mediated by glutamate transporters, caused NO generation by persistent NMDA receptor activation (188). The present study provides evidence that in the retina, glutamate dysregulation following axonal injury to RGCs leads to persistent NMDA receptor activation and NO generation. However, unlike the deleterious effects of this pathway reported earlier, our studies show that NO generation has beneficial effects in elevating levels of optic nerve regeneration. What both cases have in common is glutamate dysregulation following neuronal injury, leading to NMDA receptor dependent nitric oxide generation. The molecular trigger for the glutamate dysregulation following excitotoxic injury has not been ascertained, although a subsequent study reported blockade of neurotoxicity following a brief excitotoxic insult by the p38 kinase inhibitor SB-239063 (189). Indeed, p38 activation and Kv2.1 phosphorylation at Ser-800 have been shown to be required for translocation of the channel to the plasma membrane (102, 103) and p38 kinase is considered to be an important MAP kinase in neurodegeneration following excitotoxic and ischemic insults (190–193). Our data indicate that altered function of Kv2.1 plays an important role in the glutamate dysregulation underlying NMDA receptor activation and nitric oxide generation. Given the recent suggestion that the Kv2.1-GLT-1 interaction underlies a “neuronal injury sensor complex” (105), it is tempting to speculate that alteration of glutamate homeostasis is a critical output of this sensor.

RGC death and regenerative failure after optic nerve injury have been widely attributed to cell-autonomous changes in RGCs (6–8, 14, 15, 194–200) and to retinal micro-and macroglia (41, 201, 202). However, recent studies point to amacrine cells also playing a critical role. We previously reported an increase in an autometallographic signal in the IPL after ONC that we interpreted as an elevation of synaptic Zn^2+^ in amacrine cell terminals and that chelators of Zn^2+^ and other divalent cations improved RGC survival and axon regeneration (42, 203). Similarly beneficial effects have also been obtained by genetic deletion of the Zn^2+^ transporter ZnT3 in amacrine cells (204). In addition, Zhang et al. (43) demonstrated an increase in amacrine cell activity after ONC and found that pharmacological suppression of inhibitory neurotransmission in the retina promotes RGC survival and axon regeneration. Thus, together with the present study, increasing evidence points to amacrine cells influencing RGC survival and axon regeneration in both positive ways (via NO generation) and negative ways (through elevated synaptic inhibition). Our data also point to the involvement of bipolar cells and astrocytes in elevating extracellular glutamate that, through NMDA receptors, leads to NOS1 activation and NO generation. Finally, K^+^ efflux through Kv2.1 in RGCs may trigger this cascade that ultimately feeds back on RGCs’ ability to mount a robust regenerative response. Understanding the epistatic relationships between the cell-intrinsic changes that occur in RGCs after optic nerve injury and the changes that occur in neighboring interneurons and glia remains an important area of investiga-tion for protecting and restoring vision after traumatic, ischemic, or neurodegenerative damage to the optic nerve, with possible wider implications for understanding neurodegeneration and repair.

## Materials and Methods

### Animal use, housing, and strains

Animal studies were performed at Boston Children’s Hospital with approval of the Institutional Animal Care and Use Committee. Mice were housed in the animal facility with a 12 hr light/12 hr dark cycle (lights on from 7 AM-7 PM) and a maximum of 5 adult mice per cage. Wild-type 129S1 mice (#002448, Jackson Laboratory, ME), NOS1 conditional knock-out mice (*NOS1*^flx/flx^ from Dr. Paul Huang, MGH), Cre-driver mouse lines including Vgat-Cre (gift from Dr. Zhigang He, Boston Children’s Hospital), *VGLUT2-Cre* (#016963, Jackson Laboratory, ME), *vGLUT1-Cre* (#023527, Jackson Laboratory, ME) *Glast-CreERT* (#012586, Jackson Laboratory, ME) were used. Conditional GLT-1 knock-out mice were obtained from the founding colony at Boston Children’s Hospital (Slc1A2^tm1.1Pros^; MGI: 5752263) (82) and are available from the Mutant Mouse Resource and Research Centers Repository (B6J.129S4-*Slc1a2^tm1.1Pros^*/Mmnc; MMRRC:050600-UNC). The crossed lines *NOS1^flx/flx^ VGat-Cre*, *GLT-1^flx/flx^ VGluT1-Cre, GLT-1^flx/flx^ GLAST-CreERT were* generated in-house. Tail snips were obtained from individual animals at 3-weeks of age for genotyping by Transnetyx Inc. (TN). Only heterozygous Cre-driver mice were used for breeding and experiments. Wild-type littermates were used as controls in all experiments using genetically altered animals.

<u>Tamoxifen administration</u>: 200 mg tamoxifen (Sigma-Aldrich cat#T5648) was dissolved in 9ml corn oil (Sigma-Aldrich, cat#C-8267) with 1ml 100% ethanol to a concentration of 20 mg/ml. The solution was vortexed and left on a rocker for a few hours until completely dissolved, then stored in 4°C. Each litter of *GLT-1^flx/flx^;Glast-Cre/ERT* and control mice was given 100 µl (2 mg) via IP injection per day for 5 consecutive days at P35. Treated animals were used for experiments after P42.

### Surgical Procedures

Optic nerve surgeries and intraocular injections were carried out on male and female mice at 8-10 weeks of age (average body weight, 20-26 g) under general anesthesia as described previously (35, 42, 205). Animals were assigned to different treatment groups and, in any given experiment, surgeries were done for multiple groups at a time. Subsequent processing was performed blinded to treatment. Intraocular injection was done cautiously to avoid injury to the lens. Reagents that were injected intraocularly included an adeno-associated virus (serotype 2) expressing shRNA to knock down expression of PTEN (AAV2-shPten-mcherry) 2 weeks prior to ONC (to allow time for virally mediated knockdown of expression); Oncomodulin (Ocm, gift of Michael Henzl, Univ. Missouri) and CPT-cAMP (100 µM, Sigma Aldrich, cat# C3912) immediately after ONC to enhance levels of regeneration; NO donors diethylenetriamine NONOate (DETA-NONOate, 10 μM, Cayman Chemical, Cat# 82120) or Cay10562 (1 µM Cayman Chemical, Cat# 10010502); NOS1-selective inhibitor Nω-propyl-L-arginine (L-NPA, 30 µM, Cayman, cat# 80587), NMDA receptor antagonist D-2-amino-5-phosphonovalerate (D-AP5,1 mM, Tocris, cat# 0106), iNOS-selective inhibitor 1400W (100 µM, Cayman, cat# 81520), Superoxide Dismutase (2U/eye, Sigma-Aldrich, S5395); GLT-1 inhibitors Way213613 (10 µM, Tocris, cat# 2652) and Dihydrokainic acid (DHK, 1 mM,Tocris, cat# 0111); Kv2.1 inhibitor, RY796 (10 µM, Alomone Labs, cat# R-160) and stromatoxin (5µM, Alomone, cat# STS-350); and Cholera toxin subunit B (CTB, Sigma-Aldrich, cat# SAE0069-500UG). All agents were injected intraocularly in a volume of 3 μl, as were vehicle controls. In most experiments, agents to be tested were injected intraocularly 3 hours before ONC. SOD1 (2U) was injected on the next day, followed 20-30 minutes later by FL2 or FL2E (1 mM), and mice were perfused 1 hour later.

### NO probe preparation and NO detection

FL2 is a highly selective, membrane-permeable fluorescent NO probe synthesized in the Lippard lab as described (107). The compound was dissolved in 100% DMSO (Sigma, cat# D2650) and diluted in 1× Phosphate Buffered Saline (PBS). CuCl_2_ (20 mM in H_2_O, Sigma) was mixed 2:1 (molar ratio) to activate FL2 just before use (FL2, 200 μM). FL2E kits were purchased from Strem Chemicals (cat# 96-0396) and used according to the manufacturer’s instructions. Optimal results were obtained by intraocular injection of Cu/Zn superoxide dismutase (SOD, 2 U/eye, Sigma, cat#S5395) 20 min prior to probe injection. FL2 or FL2E was injected intravitreally (200 µM, 3.0 µl per eye) 20-30 min after the injection of SOD1 (2U), and mice were sacrificed 1 h later. At the appropriate survival times, mice were perfused transcardially with saline followed by 4% formaldehyde (FA). Eyes were dissected, post-fixed in 4% FA for 1-2 hour, cryoprotected overnight with 30% sucrose, and embedded in OCT medium. Frozen sections were cut on a cryostat at 14 μm, washed in PBS for 10 min, and mounted on treated slides. Fluorescent images were captured and relative FL2 intensity in the IPL or FL2E intensity in the GCL of the retina was analyzed using ImageJ software. Average intensities were calculated from 4-6 individual cases per condition and images are representative of these results. In most experiments, reagents to be tested were injected intraocularly 3 hours before ONC.

### Immunofluorescence

After blocking with the appropriate sera, tissue sections were incubated overnight at 4°C with primary antibodies directed to βIII tubulin (monoclonal antibody TUJ-1, 1:500, made in mouse, Covance Cat.# MMS-435P, or a rabbit polyclonal antibody, Abcam, cat# ab18207); nNOS/NOS1 (1:500, rabbit monoclonal, Cell Signaling, cat# 4231); GLT-1a (1:5000 mouse monoclonal, gift of Dr. Jeff Rothstein, Johns Hopkins University); GLT-1b (1:200 rabbit polyclonal from Dr. Niels C. Danbolt; RRID: AB_2714095), VSX2 (1:500 Sheep polyclonal; Invitrogen cat #PA1-12565), CRALBP (1:500 rabbit monoclonal, Abcam, cat# ab243664); cholera toxin B fragment (1:500, Fisher Scientific, cat# PA125635); nitrotyrosine (1:200 polyclonal, Cayman cat# 10189540); phospho-specific anti-Kv2.1 (Ser-800), from Dr. Elias Aizenman, University of Pittsburgh)(102) (1:250); total Kv2.1 (Alomone cat #APC-012)(1:500). Sections were then washed 3 times, incubated with appropriate fluorescent secondary antibodies (1:500, Molecular Probes) two hours at room temperature, and rinsed 3 times. In some cases, nuclei were counterstained with DAPI, and sections were mounted with Fluoromount G medium. Images were taken using a Nikon Air80i, Nikon E800 or Zeiss LSM700 confocal microscope.

### cGMP immunofluorescence

Retinal sections were incubated with 0.3% hydrogen peroxide in cold methanol (1:100 dilution of 30% hydrogen peroxide; Sigma Aldrich catalog # H1009), then blocked with 10% rabbit serum in Tris-buffered saline [(TBS); 50 mM Tris-HCl, pH 7.4; 0.9% NaCl] containing 1mM IBMX (Sigma Aldrich catalog # I5879) to inhibit cGMP breakdown. Sections were then incubated with sheep anti-cGMP antibody 1:1000 (obtained from Dr. Harry Steinbusch, Maastricht University) at 4°C overnight; washed 3 times with TBS next day, followed by rabbit anti-sheep biotinylated secondary antibody1:250 (Vector Laboratory catalog # BA-6000) at room temperature for 2 hours, and washing 3 times with TBS. Avidin-biotin complex (ABC) solution was prepared as specified by kit instructions (Vectastain Elite ABC-HRP system, Vector cat#PK-6100) in TBS 30 min in advance. Sections were incubated with ABC solution for 1 hour, rinsed 3 times with TBS, followed by a 25 minute incubation with DAB/peroxidase/nickel solution (DAB substrate kit, Vector catalog # SK-4100). Slides were then rinsed with TBS, then water for 15 minutes to stop the reaction, followed by dehydration using 70%, 95%, 100% ethanol then xylene. Sections were mounted with Cytoseal Mountant (Fisher Scientific cat# 22-050-262) and imaged with a Nikon E800 Air microscope.

### Visualization of protein S-nitrosylation

Protein S-nitrosylation was detected in the retina using a commercial kit (Cayman, cat# 10006525) based on the “Biotin Switch” method of Jaffrey (117), which enables detection of a fluorescent SNO bond. Tissue sections were prepared as described above for immunofluorescence and then treated according to the manufacturer’s instructions. Negative control for endogenous biotinylated proteins and labeling control were performed. Positive control slides were prepared by immersing in 1mM S-nitroso-L-glutathione (GSNO, Cayman, cat#82240) for 1 h before starting the assay (Figure1-I). Stained sections were imaged by confocal microscopy as recommended (Excitation wavelength = 490 nm, Emission = 520 nm)

### Quantitation of RGC survival and axon regeneration

RGC survival and axon regeneration were quantified as in our earlier studies (34, 42, 117, 206). Mice were euthanized at 14 days after NC and perfused transcardially with 0.9% saline and 4% paraformaldehyde (PFA; Sigma #441244) dissolved in 1X phosphate buffered saline (PBS) prepared from 10X PBS, (Bio-Rad cat# 1610780). Whole eyes and optic nerves were dissected and post-fixed in 4% PFA for two hours. Retinas were dissected and stained as whole-mounts after making four radial cuts to enable the tissue to lie flat. RGC survival was evaluated by immunostaining with an antibody to βIII-tubulin (1:500, rabbit polyclonal; Abcam, #ab18207), followed by an Alexa Fluor conjugated secondary antibody to rabbit IgG. ImageJ software was used to count βIII-tubulin-positive cells per retina from a set of fluorescently illuminated images (200x, Nikon E800) taken in 8 prespecified areas. Values were averaged to approximate RGC survival per square millimeter.

Optic nerves were transferred to 30% sucrose after post-fixation, embedded in OCT Tissue Tek Medium (Sakura Finetek/VWR), frozen, cryostat-sectioned longitudinally at 14 μm, and mounted on coated glass slides. Regenerating axons were visualized by immunostaining with a rabbit CTB antibody followed by a fluorescently labeled secondary antibody. Axons were counted manually in 4-8 longitudinal sections per case at pre-specified distances from the injury site, and these values were used to estimate the total number of regenerating axons per nerve. In the results shown in sFig. 2B, approximately half of the cases used a slightly modified version of the iDISCO tissue clearance method to visualize and quantify axon regeneration (Renier et al., PMID: 25417164; Zhang et al., PMID 39093964). Because the latter revealed somewhat more axons than our standard procedure for both negative control and experimental cases, we normalized both datasets to the respective negative control values, combined normalized data, then re-set the baseline to the average value between the two datasets.

### Autometallography (AMG)

To visualize ionic zinc we used a modified version (42) of the autometallography procedure developed by Danscher, Dyck and their colleagues in which mice are exposed to sodium selenite via i.p. injection prior to perfusion fixation (207–209). Mice were injected intraperitoneally with sodium selenite (Na_2_SeO_3_, 1.5 mg/ml in distilled H_2_O; Sigma, 15 mg/kg) and were kept alive for 4 hours to allow for optimal signal detection as described previously. An overdose of anesthesia was then given followed by transcardial perfusion with isotonic saline (Sigma) and 2.5% glutaraldehyde (GA; Ted Pella) in 0.1 M phosphate buffer (PB, pH 7.4). Whole eyes and optic nerves were dissected and immersed in 2.5% GA in 0.1 M PB fixative for 4 hours at 4°C, then transferred to 30% sucrose in PBS overnight at 4°C, embedded in OCT Tissue Tek Medium, and frozen. Tissues were cryostat-sectioned at 14 μm and mounted on glass slides (VWR). Sections were fixed in 95% ethanol, hydrated in 70% and 50% ethanol, dipped in 0.5% gelatin, and then incubated in 50 ml AMG developer solution (30 ml gum arabic, 5 ml citrate buffer, 0.43 g hydroquinone in 7.5 ml distilled water, and 0.06 g silver-lactate in 7.5 ml distilled water) for 2.5 hours at room temperature. To stop the reaction, slides were washed with running water (37°C) for 10 min. Sections were then placed in 5% sodium thiosulfate for 12 minutes, post-fixed for 30 minutes with 70% ethanol, and dehydrated with 95% and 100% ethanol followed by xylene. Five images from different areas of each retinal section were taken under brightfield illumination (600x; E800; Nikon). The intensity of the AMG signal in the IPL was analyzed using ImageJ software. AMG staining and imaging were done simultaneously for all samples to be compared with each other. Average signal intensity was calculated from 6-8 individual cases per condition. All images are representative of multiple samples.

### Optic nerve clearance

The modified iDISCO method with anterograde CTB labeling was used to visualize axon regeneration in the cleared optic nerve. After transcardial perfusion with 4% FA, optic nerves were dissected and incubated in a series of 30%, 60%, and 90% methanol in PBS for 30 minutes at each concentration, followed by dichloromethane (Millipore-Sigma, DCM, Cat. #270997)/methanol (2:1) for 20 minutes. Nerves were then transferred to dibenzyl ether (DBE, Millipore-Sigma Cat. #33630) and incubated overnight to achieve complete transparency. Cleared optic nerves were mounted using DBE for microscopy.

### Statistical Analyses

All tissue processing, quantification, and data analysis were done in a blinded fashion throughout the study. Sample sizes were based on accepted standards in the literature and prior experience from our lab. Sample size (*n*) represents total number of biological replicates in each condition across all experiments that were performed (N ≥ 2). All experiments contained positive and negative controls, with multiple experimental conditions run simultaneously. After establishing the methodology for any type of study, no cases were excluded in our data analysis. Unpaired Student’s *t*-tests, two-tailed (GraphPad Prism v.9) were used after determining normality of data distribution. Data are presented as Mean ± SEM. Difference were considered significant with *P* < 0.05 (*\*P* < 0.05, *\*\*P* < 0.01, *\*\*\*P* < 0.001).

