## Supplementary material for "An intercellular signaling pathway in the mouse retina connects Kv2.1, GLT-1, and nitric oxide synthase 1 to optic nerve regeneration": Supplemenal Figure 1 Legend

**Appendix: Supplementary Figure Legends**

**Suppl. Figure 1. NO donors partially reversed inhibitory effects of NOS1 deletion on axon regeneration.** (A) Pten deletion combined with Ocm and CPT-cAMP induced strong axon regeneration in control NOS1^wt/wt^;Vgat-Cre^+/-^ mice (*top panel*) that was diminished mice lacking NOS1 in amacrine cells (NOS1^flx/flx^ ;Vgat-Cre^+/-^, *second panel*). This loss was partially reversed by NO donors Cay10562 or Deta NONOate. (B) Quantitation of axon regeneration for conditions shown in A. n = 6-8/condition; * *P =* 0.0126, *t* = 2.858 (*KO/Cay10562 vs KO*); **P =* 0.0476, *t* = 2.229 (*KO/Nonoate vs KO*); ***P =* 0.0025, *t* = 3.807 (*VgatCre WT vs KO*); **P =* 0.0341, *t* = 2.390 (*NC vs NC/Nonoate*); ***P =* 0.0018, *t* = 4.212 (*NC vs NC/Cay10562*). In addition, intraocular application of these NO donors alone promoted minor axon regeneration after optic nerve injury (images not shown).
