## Supplementary figures and images for "An intercellular signaling pathway in the mouse retina connects Kv2.1, GLT-1, and nitric oxide synthase 1 to optic nerve regeneration"

### Supplemental Figure 1

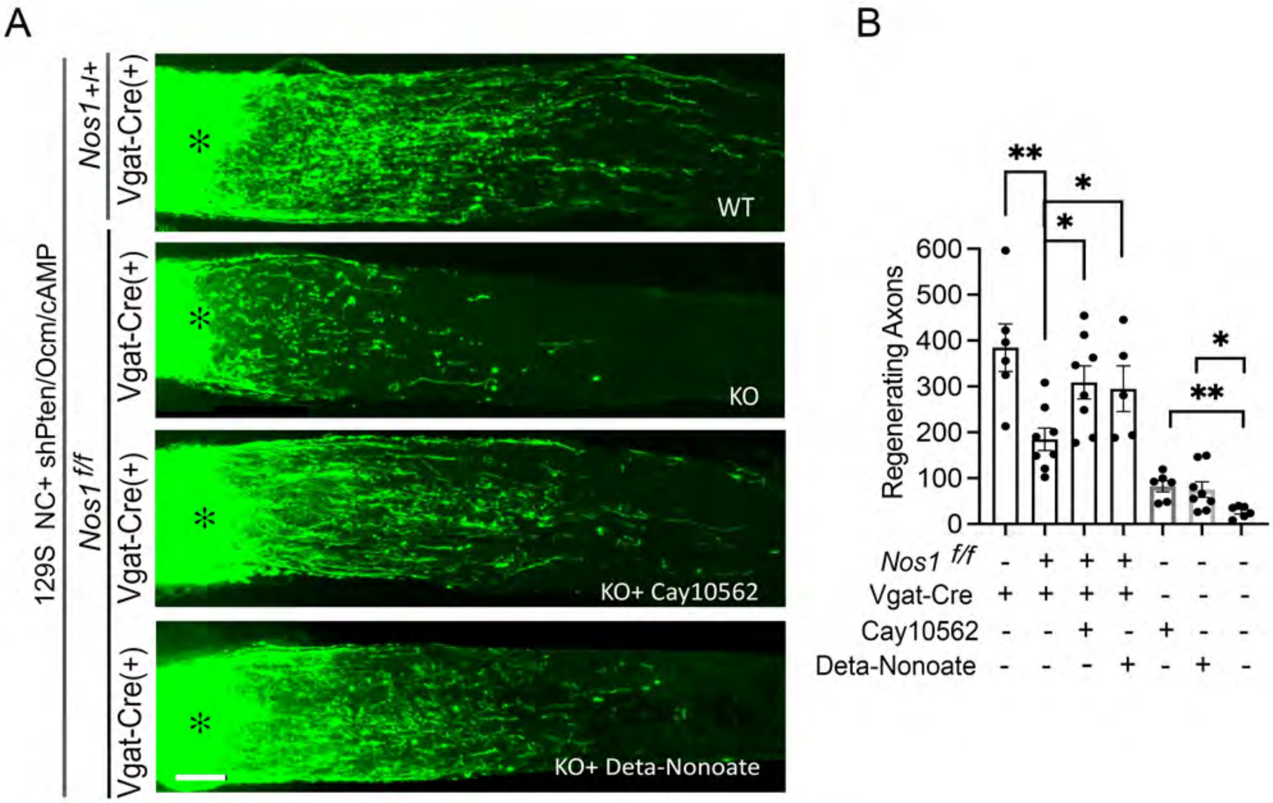
